# Self-Supervised Deep Learning Encodes High-Resolution Features of Protein Subcellular Localization

**DOI:** 10.1101/2021.03.29.437595

**Authors:** Hirofumi Kobayashi, Keith C. Cheveralls, Manuel D. Leonetti, Loic A. Royer

## Abstract

Elucidating the diversity and complexity of protein localization is essential to fully understand cellular architecture. Here, we present *cytoself*, a deep-learning approach for fully self-supervised protein localization profiling and clustering. *cytoself* leverages a self-supervised training scheme that does not require pre-existing knowledge, categories, or annotations. Training *cytoself* on images of 1,311 endogenously labeled proteins from the OpenCell database reveals a highly resolved protein localization atlas that recapitulates major scales of cellular organization, from coarse classes such as nuclear, cytoplasmic and vesicular, to the subtle localization signatures of individual protein complexes. We quantitatively validate *cytoself*’s ability to cluster proteins into organelles and protein complex clusters using a clustering score, and show that *cytoself* attains higher scores than previous unsupervised or self-supervised approaches. Finally, to better understand the inner workings of our model, we dissect the emergent features from which our clustering is derived, interpret these features in the context of the fluorescence images, and analyze the performance contributions of the different components of our approach.

---

Systematic and large-scale microscopy-based cell assays are becoming an increasingly important tool for biological discovery^1,2^, playing a key role in drug screening^3,4^, drug profiling^5,6^, and for mapping the sub-cellular localization of the proteome^7,8^. In particular, large-scale datasets based on immuno-fluorescence or endogenous fluorescent tagging comprehensively capture localization patterns across the human^9,10^ and yeast proteome^11^. Together with recent advances in computer vision and deep learning^12^, such datasets are poised to help systematically map the cell’s spatial architecture. This situation is reminiscent of the early days of genomics, when the advent of high-throughput and high-fidelity sequencing technologies was accompanied by the development of novel algorithms to analyze, compare, and categorize these sequences, and the genes therein. However, images pose unique obstacles to analysis. While sequences can be compared against a frame of reference (i.e. genomes), there are no such references for microscopy images. Indeed, cells exhibit a wide variety of shapes and appearances that reflect a plurality of states. This rich diversity is much harder to model and analyze than, for example, sequence variability. Moreover, much of this diversity is stochastic, posing the additional challenge of separating information of biological relevance from irrelevant variance. The fundamental computational challenge posed by image-based screens is therefore to extract well-referenced vectorial representations that faithfully capture only the relevant biological information and allow for quantitative comparison, categorization, and biological interpretation of protein localization patterns.

Previous approaches to classify and compare images have relied on engineered features that quantify different aspects of image content – such as cell size, shape and texture^13–16^. While these features are, by design, relevant and interpretable, the underlying assumption is that all the relevant features needed to analyze an image can be identified and appropriately quantified. This assumption has been challenged by deep learning’s recent successes^17^. On a wide range of computer vision tasks such as image classification, hand-designed features cannot compete against learned features that are automatically discovered from the data itself^18,19^. Assuming features are available, the typical approach consists of boot-strapping the annotation process by either (i) unsupervised clustering techniques^20,21^, or (ii) manual curation and supervised learning^22,23^. In the case of supervised approaches, human annotators examine images and assign annotations, and once sufficient data is garnered, a machine learning model is trained in a supervised manner, and later applied to unannotated data^17,18,23,24^. Another approach consists of reusing models trained on natural images to learn generic features upon which supervised training can be bootstrapped^5,25,26^. While successful, these approaches suffer from potential biases, as manual annotation imposes our own preconceptions. Overall, the ideal algorithm should not rely on human knowledge or judgments, but instead automatically synthesize features and analyze images without a priori assumptions – that is, solely on the basis of the images themselves.

Recent advances in computer vision and machine learning have shown that forgoing manual labeling is possible and nears the performance of supervised approaches^27,28^. Instead of annotating datasets, which is inherently non-scalable and labor-intensive, self-supervised models can be trained from large uncurated datasets^11,29–32^. Self-supervised models are trained by formulating an auxiliary *pretext task*, typically one that withholds parts of the data and instructs the model to predict them^33^. This works because the task-relevant information within a piece of data is often distributed over multiple observed dimensions^30^. For example, given the picture of a car, we can recognize the presence of a vehicle even if many pixels are hidden, perhaps even when half of the image is occluded. Now, consider a large dataset of pictures of real-world objects (e.g. ImageNet^34^). Training a model to predict missing parts from these images forces it to identify their important features^32^. Once trained, the vectorial *representations* that emerge from pretext tasks capture the important features of the images, and can be used for comparison and categorization^35^.

Here, we present the development, validation and utility of *cytoself*, a deep learning-based approach for fully selfsupervised protein localization profiling and clustering. The key innovation is a pretext task that ensures that the localization features that emerge from different images of the same protein are helpful to distinguish its images from the images of other proteins in the dataset. We demonstrate the ability of *cytoself* to reduce images to feature profiles characteristic of protein localization, validate their utility to predict protein assignment to organelles and protein complexes, and compare with previous image featurization approaches.

## Results

### A robust and comprehensive image dataset

A prerequisite to our deep-learning approach is a collection of high-quality images of fluorescently tagged proteins obtained under uniform conditions. Our OpenCell^10^ dataset of live-cell confocal images of 1,311 endogenously tagged proteins (opencell.czbiohub.org) meets this purpose. We reasoned that providing a fiducial channel could provide a useful reference frame for our model to capture protein localization. Hence, in addition to imaging the endogenous tag (split mNeonGreen2), we also imaged a nuclear fiducial marker (Hoechst 33342) and converted it into a distance map (see Methods). On average, we imaged the localization of a given protein in about 18.59 field of view (FOV). Approximately 45 cropped images from each FOV containing 1-3 cells were then extracted for a total of 800 cropped images per protein. This scale, as well as the uniform conditions under which the images were collected, were important because our model must learn to ignore image variance and instead focus on protein localization. Finally, in our approach all images that represent the same protein were labeled by the same unique identifier (we used the corresponding synthetic cell line identifier, but the identifier can be arbitrary). This identifier does not carry any explicit localization information, nor is it linked to any metadata or annotations, but rather is used to link together all the different images of the same protein.

### A Deep Learning model to generate vectorial image representations

Our deep learning model is based on the Vector Quantized Variational Autoencoder architecture (VQ- VAE^37,38^). In a classical VQ-VAE, images are encoded into a quantized *latent* representation, a vector, and then decoded to reconstruct the input image (see Fig. 1). The encoder and decoder are trained so as to minimize distortion between input and output images. The representation produced by the encoder is assembled by arraying a finite number of *symbols* (indices) that stand for vectors in a *codebook* (Fig. 1b, Supp. Fig.1). The codebook vectors themselves evolve during training so as to be most effective for the encoding-decoding task^37^. The latest incarnation of this architecture (VQ-VAE-2^36^) introduces a hierarchy of representations that operate at multiple spatial scales (termed VQ1 and VQ2 in the original VQ-VAE-2 study). We chose this architecture as a starting point because of the large body of evidence that suggests that quantized architectures currently learn the best image representations^37,38^. As shown in Fig. 1b we developed a variant that utilizes a split vector quantization scheme to improve quantization at large spatial scales (see Methods, Supp. Fig. 1). This new approach to vector quantization achieves better perplexity as shown in Fig. 1c which means better codebook utilization.

**Figure 1:**
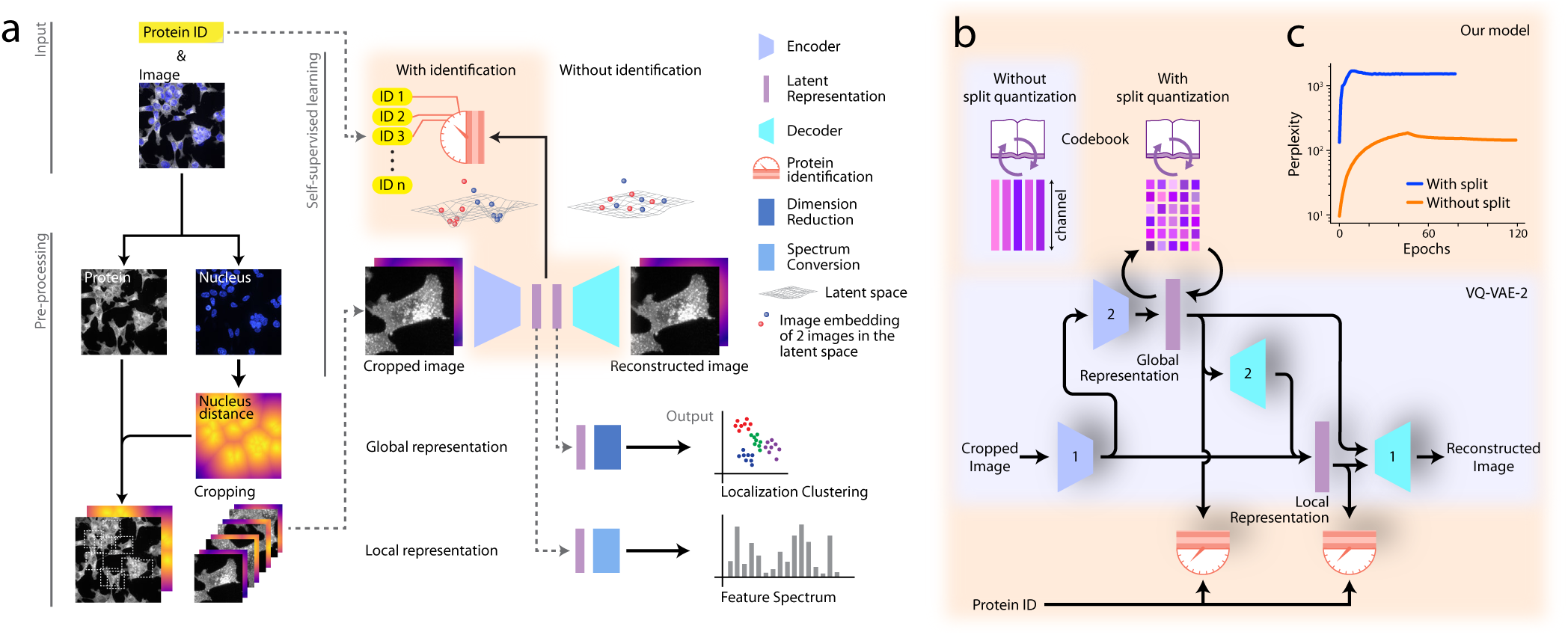
Self-supervised deep learning of protein subcellular localization with *cytoself*. **(a)** Workflow of the learning process. Only images and the proteins identifiers are required as input. We trained our model with a second fiducial channel for the cell nuclei, but its presence is optional as its performance contribution is negligible (see Fig. 4). The protein identification pretext task ensures that images corresponding to the same or similar proteins have similar representations. **(b)** Architecture of our VQ-VAE-2^36^ based Deep Learning model featuring our two innovations: split-quantization and protein identification pretext task. Numbers in the encoders and decoders indicate *encoder1, encoder2, decoder1* or *decoder2* (Supp. File. 5). Global Representation and Local Representation use different codebooks. **(c)** The level of utilization of the codebook (i.e. perplexity) increases and then saturates during training and is enhanced by applying split quantization.

### Protein localization encoding via self-supervision

Our model consists of two pretext tasks applied to each individual cropped image: First, it is tasked to encode and then decode the image as in the original VQ-VAE model. Second, it is tasked to predict the protein identifier associated with the image solely on the basis of the encoded representation. In other words, that second task aims to predict, for each single cropped image, which one of the 1,311 proteins in our library the image corresponds to. The first task forces our model to distill lower-dimensional representations of the images, while the second task forces these representations to be strong predictors of protein identity, This second task assumes that protein localization is the primary image information that is correlated to protein identity. Therefore, predicting the identifier associated with each image is key to encouraging our model to learn localization-specific representations. Interestingly, it is acceptable, and in some cases perfectly reasonable, for these tasks to fail. For example, when two proteins have identical localization, it is impossible to resolve the identity of the tagged proteins from images alone. Moreover, the autoencoder might be unable to perfectly reconstruct an image from the intermediate representation, when constrained to make that representation maximally predictive of protein identity. It follows that the real output of our model is not the reconstructed image, nor the predicted identity of the tagged protein, but instead the distilled image representations, which we refer to as ‘localization encodings’ obtained as a necessary byproduct of satisfying both pretext tasks. Specifically, our model encodes two representations for each image that correspond to two different spatial scales: the local and global representations, that correspond to VQ1 and VQ2 respectively. The global representation captures large-scale image structure scaled-down 4 × 4 pixel image with 576 features (values) per pixel. The local representation captures finer spatially resolved details (25 × 25 pixel image with 64 features per pixel). We use the global representations to perform localization clustering, and the local representations to provide a finer and spatially resolved decomposition of protein localization.

### Mapping the protein localization landscape with *cytoself*

Obtaining image representations that are highly correlated with protein localization and invariant to other sources of heterogeneity (i.e. cell state, density, and shape) is only the first step for biological interpretation. Indeed, while these representations are lower dimensional than the images themselves, they still have too many dimensions for direct inspection and visualization. Therefore, we performed dimensionality reduction using the Uniform Manifold Approximation and Projection (UMAP) algorithm on the set of global localization-encodings obtained from all images (see Methods). In the resulting UMAP (Fig. 2) each point represents a single (cropped) image in our test dataset (i.e. 10% of entire dataset, see Methods) which collectively form a highly detailed map of the full diversity of protein localizations. This protein localization atlas reveals an organization of clusters and sub-clusters reflective of eukaryotic subcellular architecture. We can evaluate and explore this map by labeling each protein according to its subcellular localization obtained from independent manual annotations of our image dataset (Supp. File 2). The most pronounced delineation corresponds to nuclear (top right) versus non-nuclear (bottom left) localizations (encircled and expanded in Fig. 2, top right and bottom left, respectively). Within the nuclear cluster, sub-clusters are resolved that correspond to nucleoplasm, chromatin, nuclear membrane, and the nucleolus. Strikingly, within each region, tight clusters that correspond to specific cellular functions can be resolved (dashed outlines). For example, subunits involved in splicing (SF3 splicesome), transcription (core RNA polymerase) or nuclear import (Nuclear pore) cluster tightly together (outlined in Fig. 2, dashed outlines). Similarly, sub-domains emerge within the non-nuclear cluster, the largest corresponding to cytoplasmic and vesicular localizations. Within these domains are several very tight clusters corresponding to mitochondria, ER exit sites (COPII), ribosomes, and clathrin coated vesicles (Fig. 2). The many gray dots outside of these discrete localization domains, often correspond to proteins that exhibit mixed localization patterns (Fig. 2). Prominent among these is a band of proteins interspersed between the nuclear and non-nuclear regions (expanded in Fig. 3a). Representative proteins chosen along that path show a continuous gradation from mostly cytoplasmic to mostly nuclear localization.

**Figure 2:**
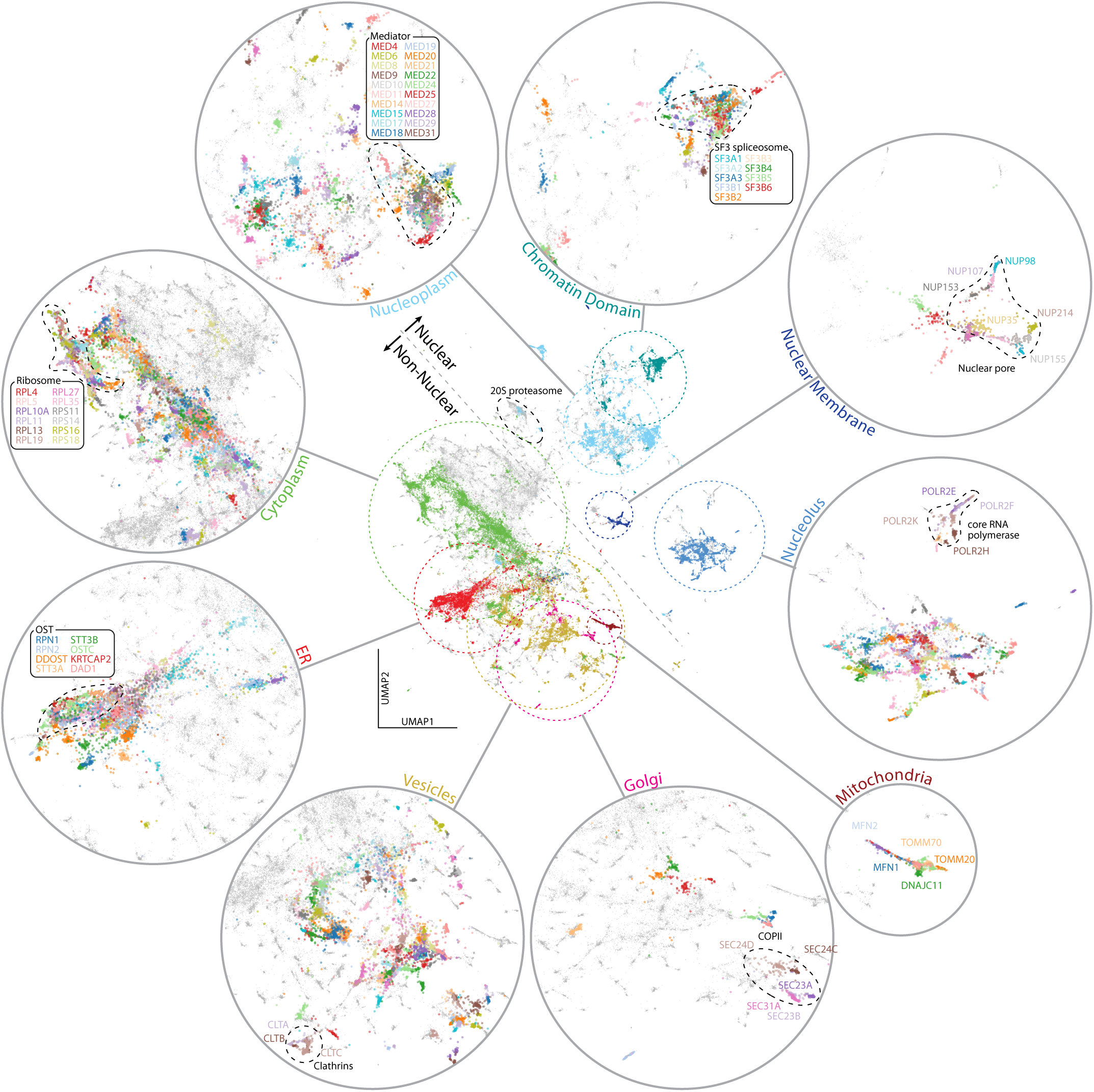
High-resolution Protein Localization Atlas. Each point corresponds to a single image from our test dataset of 109,751 images. To reveal the underlying structure of our map, each point in the central UMAP is colored according to 11 distinct protein localization categories (mitochondria, vesicules, nucleoplasm, cytoplasm, nuclear membrane, ER, nucleolus, Golgi, chromatin domain). These categories are expanded in the surrounding circles. Tight clusters corresponding to functionally-defined protein complexes can be identified within each localization category. Only proteins with a clear and exclusive localization pattern are colored, gray points correspond to proteins with other or mixed localizations. Within each localization category, the resolution of *cytoself* representations is further illustrated by labeling the images corresponding to individual proteins in different colors (dashed circular inserts). Note that while the colors in the central UMAP represent different cellular territories, colors in the inserts are only used to delineate individual proteins, and do not correspond to the colors used in the main UMAP. The list of annotated proteins are indicated in the Supp. File 3

**Figure 3:**
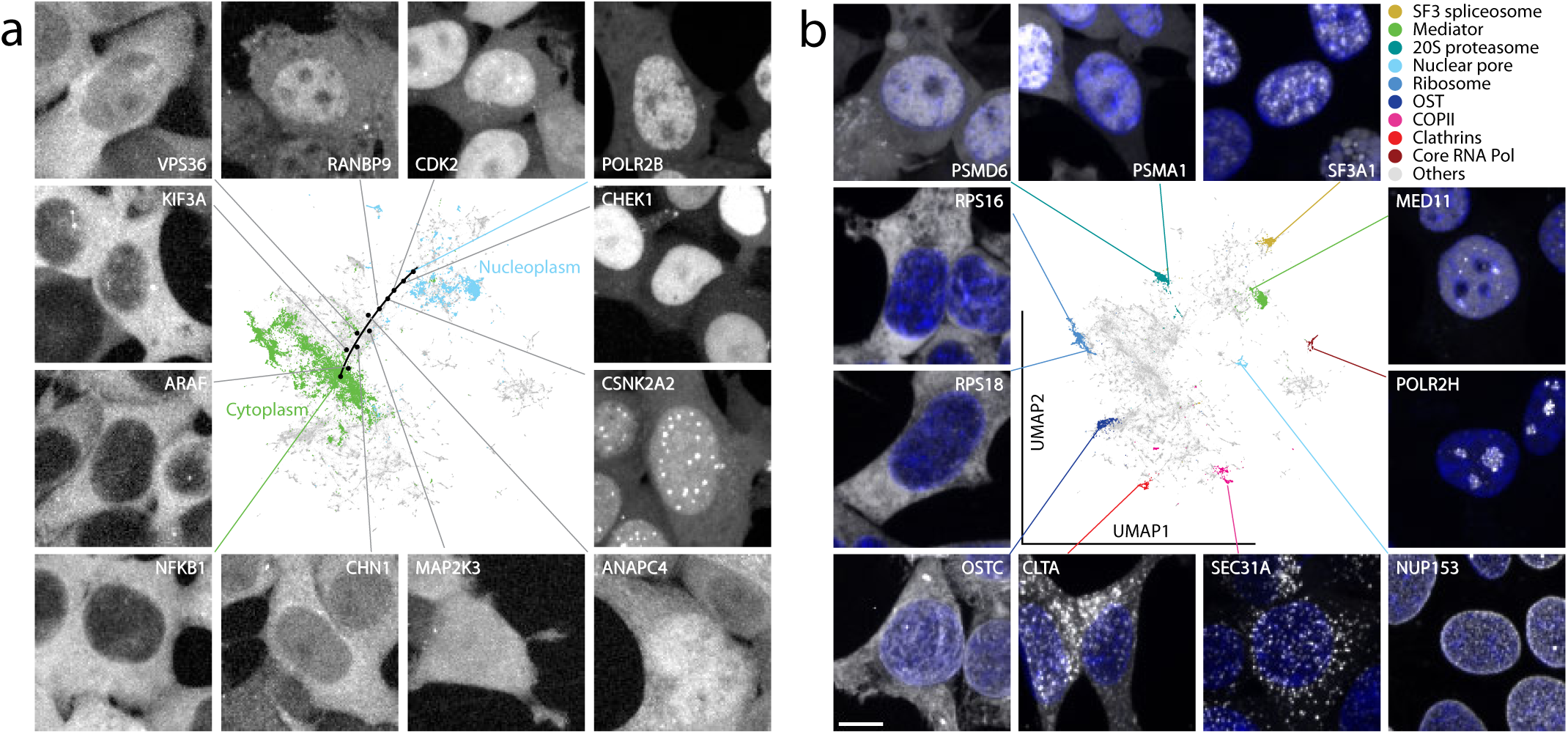
Exploring the Protein Localization Atlas. **(a)** Representative images of proteins localized along an exemplary path across the nuclear-cytoplasmic transition and over the ‘grey’ space of mixed localisations. **(b)** The subunits of well-known and stable protein complexes tightly cluster together. Moreover, the complexes themselves are placed in their correct cellular contexts. Different proteins have different expression levels, hence we adjust the brightness of each panel so as to make all localizations present in each image more visible (only min max intensities are adjusted, no gamma adjustment used). All representative images were randomly selected. Protein localization is displayed in gray color in both panels, nuclei in panel **(b)** is displayed in blue. Scale bars, 10 *µm*.

### Quantifying *cytoself*’s clustering performance

To validate our results, clustering scores were computed (see Methods, Fig. 4, and Supp. Table 1) using two ground-truth annotation datasets to capture known protein localization at two different scales: the first is a manually curated list of proteins with unique organelle-level localizations (Supp. File 3), whereas the second is a list of proteins participating in stable protein complexes derived from the CORUM database^39^ (Supp. File 1). While the first ground-truth dataset helps us assess how well our encodings cluster together proteins belonging to the same organelles, the second helps us assess whether proteins interacting within the same complex – and thus functionally related – are in proximity.

**Figure 4:**
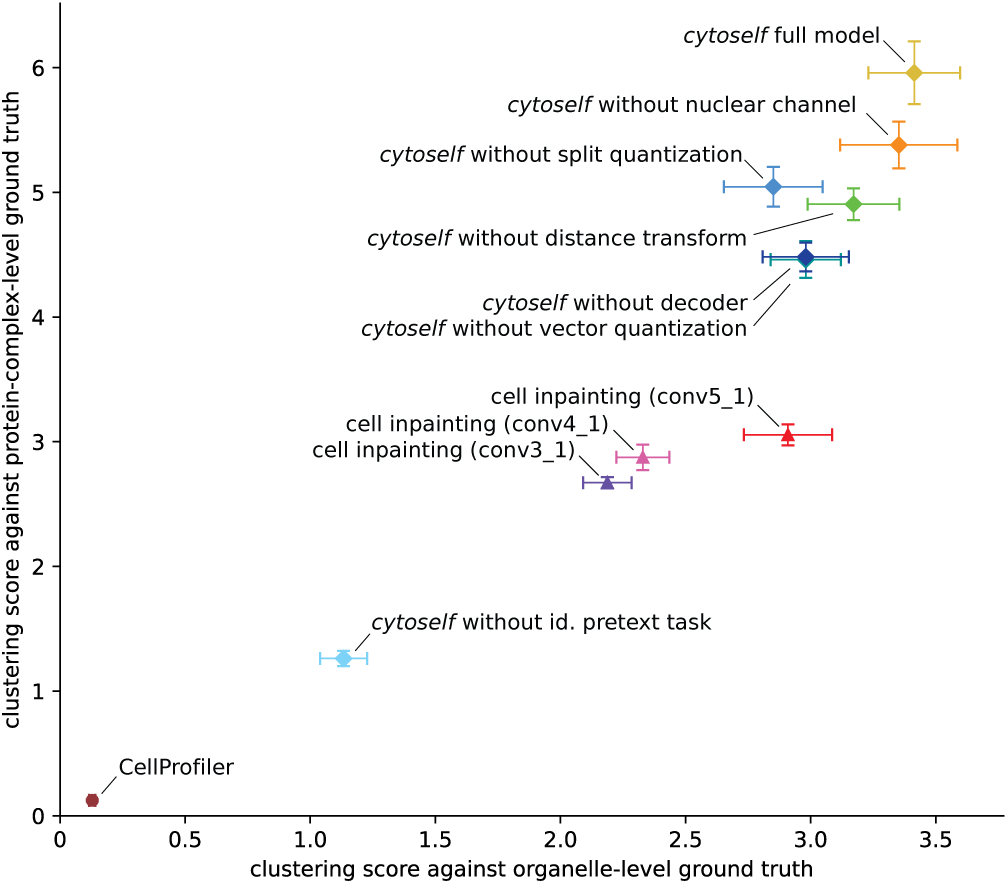
Clustering performance quantifies the effect of removing the indicated components of our model on its performance. For each model variation, we trained five model instances, compute UMAPs for ten random seeds, compute clustering scores using organelle-level and protein-complex-level ground truth, and then report mean and standard error of the mean.

### Identifying *cytoself*’s essential components

To evaluate the impact of different aspects of our model on its clustering performance, we conducted an *ablation study*. We retrained our model and recomputed a protein localization UMAPs after individually removing each component or input of our model (see Supp. Fig. 4, 5), including: (a) the nuclear fiducial channel, (b) the distance transform applied to nuclear fiducial channel, (c) the split vector quantization, and (d) the identification pretext task. We also quantitatively evaluated the effects of their ablation by computing clustering scores for different variants (Fig. 4 and Supp. Table 1). The UMAP results and scores from both sets of ground-truth labels make it clear that the single most important component of *cytoself*, in terms of clustering performance, is the protein identification pretext task. The remaining components – the nuclear channel, split quantization, vector quantization, etc – are important but not crucial. Interestingly, forgoing the fiducial nuclear channel entirely led to the smallest decrease in clustering score, suggesting that our approach works well even in the absence of any fiducial marker – a notable advantage that widens the applicability of our approach and greatly simplifies the experimental design^40^. Overall, our data shows a robust fit with ground truth. In conclusion, although all features contribute to the overall performance of our model, the identification pretext task is the key and necessary ingredient.

### Comparative performance of *cytoself*

Other unsupervised (CellProfiler^14^) or self-supervised (Cell inpainting^11^) approaches for image featurization have been previously developed. We therefore applied these methods to the OpenCell image dataset and then compared the results to that obtained by *cytoself*. UMAPs were calculated for each model (see Methods) and compared with our set of ground-truth organelles and protein complexes. As can be seen (Supp. Fig. 6, 7 and 14), the resolution obtained by *cytoself* exceeded that of both previous approaches. This was also apparent in our calculations of clustering scores (see Fig. 4 and Supp. Table 1).

### Revealing subtle protein localization differences not annotated in existing image-based localization databases

The key advantage of self-supervised approaches is that they are not limited by the quality, completeness or granularity of human annotations. To demonstrate this, we asked whether *cytoself* could resolve subtle localization differences that are not present in image-derived manual annotations – focusing on proteins localized to intracellular vesicles. Even though several known sub-categories of vesicles exist (e.g. lysosomes versus endosomes), in both OpenCell and HPA (Human Protein Atlas) annotations, these groups are annotated simply as ‘vesicles’. This reflects the difficulty for human curators to accurately distinguish and classify localization sub-categories that present similarly in the images. To test whether our self-supervised approach manages to capture these sub-categories, we focused on a curated list of endosomal as well as lysosomal proteins identified by an objective criterion. Specifically, we selected proteins annotated as lysosomal (GO:000576500) or endosomal (GO:0031901) in Uniprot^41^ (excluding targets annotated to reside in both compartments), and for which localization in each compartment has been confirmed independently by mass spectrometry^42,43^. As shown in Supp. Fig. 12, the representation of the lysosomal versus endosomal images derived from *cytoself* form two distinct, well-separated clusters (*p <* 10^−3^, Mann–Whitney U test). This demonstrates that self-supervised approaches are not limited by ground truth annotations and can reveal subtle differences in protein localization not explicitly present in existing databases.

### Extracting feature spectra for quantitative analysis of protein localization

*cytoself* can generate a highly resolved map of protein localization on the basis of distilled image representations. Can we dissect and understand the features that make up these representations and interpret their meaning? To identify and better define the features that make up these representations, we created a *feature spectrum* of the main components contributing to each protein’s localization encoding. The spectra were constructed by calculating the histogram of codebook feature indices used in each image (see Supp. Fig. 2, and Fig. 1a, and Methods for details). To group related and possibly redundant features together, we performed hierarchical biclustering^44^ (Fig. 5a), and thus obtained a meaningful linear ordering of features by which the spectra can be sorted. This analysis reveals feature clusters of which we manually select 11 from the top levels of the feature hierarchy (Fig. 5a, bottom). Representative images from each cluster illustrate the variety of distinctive localization patterns that are present at different levels across all proteins. For example, the features in the first clusters (*i, ii, iii*, and *iv*) corresponds to a wide range of diffuse cytoplasmic localizations. Cluster *v* features are unique to nucleolus proteins. Features making up cluster *vi* correspond to very small and bright punctate structures, that are often characteristic of centrosomes, vesicules, or cytoplasmic condensates. Clusters *vii, viii*, and *x* correspond to different types of nuclear localization patterns. Cluster *ix* are dark features corresponding to non-fluorescent background regions. Finally, cluster *xi* corresponds to a large variety of more abundant, punctate structures occurring throughout the cells, primarily vesicular, but also Golgi, mitochondria, cytoskeleton, and subdomains of the ER. For a quantitative evaluation, we computed the average feature spectrum for all proteins belonging to each localization category present in our reference set of manual annotations (e.g., Golgi, nucleolus, etc., see Fig. 5b and Supp. File. 2). This analysis confirms that certain spectral clusters are specific to certain localization categories and thus correspond to characteristic textures and patterns in the images. For example, the highly specific chromatin and mitochondrial localizations both appear to elicit very narrow responses in their feature spectra.

**Figure 5:**
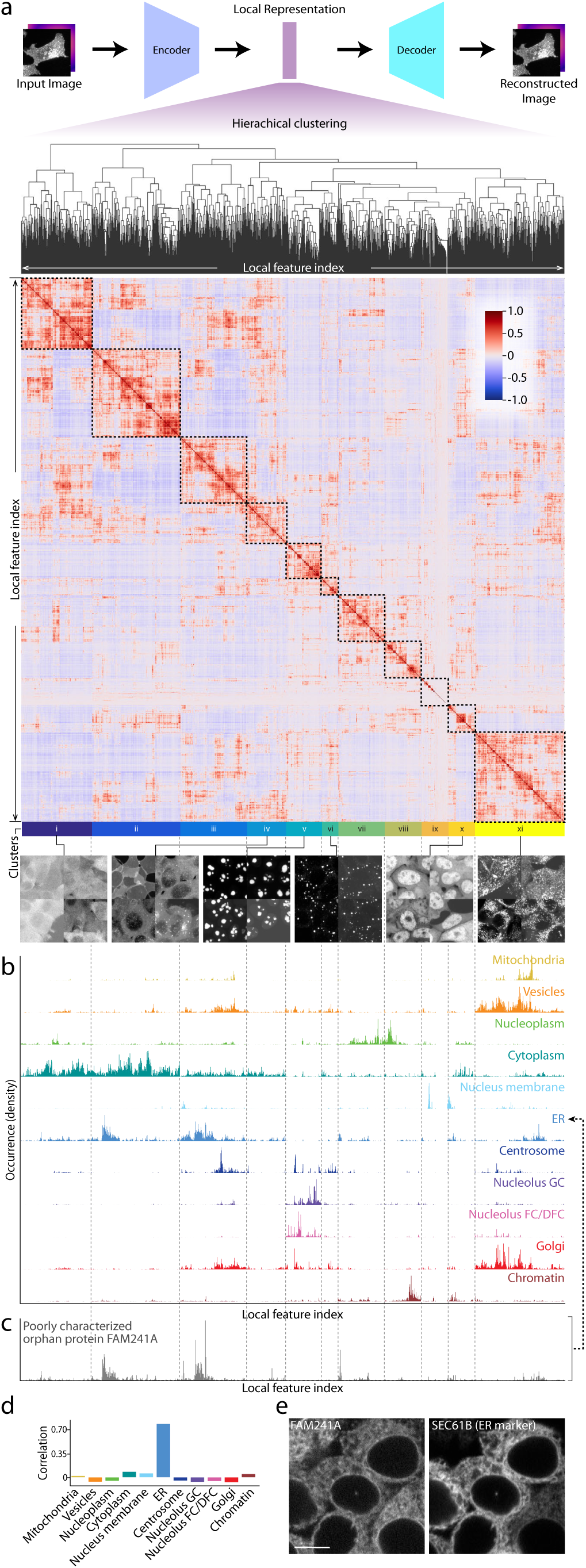
(Continued) **(a)** Features in the local representation are reordered by hierarchical clustering to form a feature spectra (see Supp. Fig. 2). The color bar indicates the strength of correlation. Negative values indicate anti-correlation. On the basis of the feature clustering, we manually identified 11 primary top-level clusters, which are illustrated with representative images (see also Supp. Fig. 3). **(b)** Average feature spectrum for each unique localization family. Occurrence indicates how many times a quantized vector is found in the local representation of an image. All spectra, as well as the heatmap are vertically aligned. **(c)** The feature spectrum of FAM241A, a poorly characterized orphan protein. **(d)** Correlation between FAM241A and other unique localization categories. The highest correlation is 0.777 with ER, next is 0.08 with cytoplasm. **(e)** Experimental confirmation of the ER localization of FAM241A. The localization of FAM241A to the ER is experimentally confirmed by co-expression of a classical ER marker (mCherry fused to the SEC61B transmembrane domain, left) in FAM241A-mNeonGreen endogenously tagged cells (right). The ER marker is expressed using transient transfection. As a consequence, not all cells are transfected and levels of expression may vary. Scale bar: 10*µm*

### Predicting protein organelle localization with *cytoself*

We next asked whether feature spectra could be used to predict the localizations of proteins not present in our training data. For this purposes, we computed the feature spectrum of FAM241A – a protein of unknown function that was not present in the training dataset. Its spectrum is most correlated to the consensus spectrum of proteins belonging to the endoplasmic reticulum (see Fig. 5b-d and Supp. Fig.8). Indeed, FAM241A’s localization to the ER is validated experimentally by co-expression experiments showing that endogenously tagged FAM241A colocalizes with an ER marker (Supp. Fig. 5e). In a companion study^10^, we further validated by mass-spectrometry that FAM241A is in fact a new subunit of the OST (oligosaccharyltransferase) complex, responsible for co-translational glycosylation at the ER membrane. Our successful prediction of the localization of FAM241A suggests that *cytoself* encodings can be used more generally to predict organelle-level localization categories. To demonstrate this, we focused on proteins annotated to localize to a single organelle (i.e., not multi-localizing, see Supp. File 2). For each of these proteins, we re-computed the representative spectra for each of their known localization categories (i.e. ER, mitochondria, Golgi, etc.), but leaving out that protein, and then applied the same spectral correlation as described for FAM241A. This allows us to predict the protein’s localization by identifying the organelle with which its spectrum correlates best. Supp. Fig. 9 shows the accuracy of the predictions derived from this approach: for 88% of proteins, the spectra correlate best with the correctly annotated organelle. For 96% of proteins, the correct annotation is within the top 2 predictions, and for 99% it is within the top 3 predictions. Overall, this form of cross-validation verifies the discriminating power of our spectra and shows that the information encoded in each protein’s spectrum can be interpreted to predict subcellular localization.

### *cytoself* applicability beyond OpenCell data

Finally, we asked whether *cytoself* can make reasonable protein localization predictions on images from datasets other than OpenCell. To answer this question we chose data from the Allen Institute Cell collection^45^, which also uses endogenous tagging and live-cell imaging, making their image data directly comparable to ours. Importantly, the Allen collection uses a cell line (WTC11, iPSC) whose overall morphology is very different from the cell line used for OpenCell (HEK293T). We reasoned that if *cytoself* manages to capture true features of protein localization, a compelling validation would be that its performance would be cell-type agnostic. Indeed, localization encodings for images from the Allen dataset generated by a *cytoself* model trained only on OpenCell images revealed strong concordance between the embeddings of the same (or closely related) protein that were imaged in both cell datasets (see Supp. Fig. 11b). This shows that our model manages to predict protein localization even under conditions that were not directly included for training. To facilitate comparison we focused on the intersection set of nine proteins found in both the OpenCell and Allen datasets (Supp. Fig. 11a). We ran the same organelle localization prediction task and observed that in 88% (8 out of 9) of cases the correct localization is among the top 3 predictions (Supp. Fig. 10).

### Hypothesizing protein complex membership from images

The resolving power of our approach is further illustrated by examining known stable protein complexes, which are found to form well delineated clusters in our localization UMAP (see examples highlighted in Fig. 2, dashed line). Fluorescent images of 11 representative subunits from these complexes illustrate these discrete localization patterns (Fig. 3b). To substantiate these observations quantitatively, we computed the correlation of feature spectra between any two pairs of proteins in our dataset. This showed a significantly higher correlation for protein pairs annotated to belong to the same complex in CORUM compared to pairs that are not (*p <* 10^−10^, Mann-Whitney U Test; Supp. Fig. 15a). To further evaluate the relationship between proximity in feature space and protein complex membership, we examine the proportion of proteins in OpenCell that share complex membership with their most-correlated neighboring protein (see Supp. Fig. 15b). We find that 83% of highly correlated (*>* 0.95) neighbor proteins are in the same complex, and even more weakly correlated (*>* 0.8) proteins are localized to complexes 60% of the time. These results confirm that close proximity in feature space is highly indicative of protein complex membership and suggests that the features derived by *cytoself* contain fine-grained information related to very specific functional relationships.

## Discussion

We have shown that a self-supervised training scheme can produce image representations that capture the organization of protein subcellular localization (Fig. 2), solely on the basis of a large dataset of fluorescence images. Our model generates a high-resolution localization atlas capable of delineating not only organelles, but also some protein complexes. Moreover, we can represent each image with a feature spectrum to better analyze the repertoire of localization patterns present in our data. Since a protein’s localization is highly correlated with its cellular function, *cytoself* will be an invaluable tool to make preliminary functional predictions for unknown or poorly studied proteins, and for quantitatively studying the effect of cellular perturbations and cell state changes on protein subcellular localization.

Our method makes few assumptions, but imposes two pretext tasks. Of these, requiring the model to identify proteins based solely on their localization encodings was essential. We also included Hoescht DNA-staining as a fiducial marker, assuming that this would provide a spatial reference frame against which to interpret localization. Surprisingly however, this added little to the performance of our model in terms of clustering score. By comparison, the self-supervised approach by Lu *et al*.^11^ applied a pretext task that predicts the fluorescence signal of a labeled protein in one cell from its fiducial markers and from the fluorescence signal in a second, different cell from the same field of view. This assumes that fiducial channels are available, and that protein fluorescence is always well-correlated to these fiducials. In contrast, our approach only requires a single fluorescence channel and yields better clustering performance (Supp. Fig.6, 7, Supp. Table1).

The main difference between our work and the problem addressed by the Human Protein Atlas Image Classification competition^23^ is that we do not aim to predict localization patterns on the basis of manual annotations. Instead, we aim to discover de-novo the landscape of possible protein localizations. This frees us from the limitations of these annotations which include: lack of uniform coverage, uneven annotation granularity, human perceptive biases, and existing preconceptions on the architecture of the cell. This also circumvents the time-intensive efforts required to manually annotate images.

While powerful, there remains a few avenues for further development of *cytoself*. For example, we trained our model using two-dimensional maximum-intensity z-projections and have not yet leveraged the full 3D confocal images available in the OpenCell^10^ dataset. The third dimension might confer an advantage for specific protein localization patterns that are characterized by specific variations along the basal-apical cell axis. Other important topics to explore are the automatic suppression of residual batch effects, improved cell segmentation via additional fiducial channels, use of label-free imaging modalities, as well as automatic rejection of anomalous or uncharacteristic cells from our training dataset. More fundamentally, significant conceptual improvements will require an improved self-supervised model that explicitly disentangles cellular heterogeneity from localization diversity^46^.

More generally, our ability to generate data is outpacing the human ability to manually annotate it. Moreover, there is already ample evidence that abundance of image data has a *quality all its own*, i.e. increasing the size of an image dataset often has higher impact on performance than improving the algorithm itself^47^. We envision that self-supervision will be a powerful tool to handle the large amount of data produced by novel instruments, end-to-end automation, and high-throughput image-based assays.

## Methods

### Fluorescence image dataset

All experimental and imaging details can be found in our companion study^10^. Briefly, HEK293T cells were genetically tagged with split-fluorescent proteins (FP) using CRISPR-based techniques^48^. After nuclear staining with Hoechst 33342, live cells were imaged with a spinning-disk confocal microscope (Andor Dragonfly). Typically, 18 fields of view were acquired for each one of the 1,311 tagged protein, for a total of 24,382 three-dimensional images of dimension 1024 × 1024 × 22 voxel.

### Image data pre-processing

Each 3D confocal image was first reduced to two dimensions using a maximum-intensity projection along the z-axis followed by downsampling in the XY dimensions by a factor of two to obtain a single 2D image per field of view (512 × 512 pixel). To help our model make use of the nuclear fiducial label we applied a distance transform to a nucleus segmentation mask (see below). The distance transform is constructed so that pixels within the nucleus were assigned a positive value that represents the shortest distance from the pixel to the nuclear boundary, and pixel values outside of the nucleus were assigned a negative value that represents the shortest distance to the nuclear boundary (see Fig. 1a). For each dual-channel and full field-of-view image, multiple regions of dimension 100 × 100 pixel were computationally chosen so that at least one cell is present and centered, resulting in a total of 1,100,253 cropped images. Cells (and their nuclei) that are too close to image edges are ignored. The raw pixel intensities in the fluorescence channel are normalized between 0 and 1, and the nuclear distance channel is normalized between −1 and 1.

### Nucleus segmentation

Nuclei are segmented by first thresholding the nucleus channel (Hoechst staining) and then applying a custom iterative refinement algorithm to eliminate under segmentation of adjacent nuclei. In the thresholding step, a low-pass Gaussian filter is first applied, followed by intensity thresholding using a threshold value calculated by Li’s iterative Minimum Cross Entropy method^49,50^. The resulting segmentation is refined by applying the following steps: (i) we generate a ‘refined’ background mask by thresholding the laplace transform at zero, (ii) we morphologically close this mask and fill holes to eliminate intra-nuclear holes or gaps (empirically, this requires a closing disk of radius at least 4 pixels), (iii) we multiply this ‘refined’ mask by the existing background mask to restore any ‘true’ holes/gaps that were present in the background mask, (iv) we generate a mask of local minima in the laplace transform, using an empirically-selected percentile threshold, and finally (v) we iterate over regions in this local-minima mask and remove them from the refined mask if they partially overlap with the background of the refined mask.

### Detailed model architecture

All details of our model architecture are given in Suppl. File. 5 and diagrammed in Fig. 1b. First, the input image (100 × 100 × 2 pixel) is fed to *encoder1* to produce a set of latent vectors which have two destinations: *encoder2* and *VQ1 VectorQuantizer* layer. In the *encoder2*, higher level representations are distilled from these latent vectors and passed to the output. The output of *encoder2* is quantized in the *VQ2 VectorQuantizer* layer to form what we call “global representation”. The global representation is then passed to the *fc2* classifier for purposes of the classification pretext task. It is also passed on to *decoder2* to reconstruct the input data of *encoder2*. In this way, *encoder2* and *decoder2* form an independent autoencoder. The function of layer *mselyr1* is to adapt the output of *decoder2* to match the dimensions of the output of *encoder1*, which is identical to the dimensions of the input of *encoder2*. In the case of the *VQ1 VectorQuantizer* layer, vectors are quantized to form what we call the local representations. The local representation is then passed to the *fc1* classifier for purposes of the classification pretext task, as well as concatenated to the global representation that is resized to match the local representations’ dimensions. The concatenated result is then passed to the *decoder1* to reconstruct the input image. Here, *encoder1* and *decoder1* form another autoencoder.

### Split quantization

In the case of our global representation, we observed that the high level of spatial pooling required (4×4 pixel) led to codebook under-utilization because the quantized vectors are too few and each one of them has too many dimensions (Fig. 1b). To solve this challenge we introduced the concept of *split quantization*. Instead of quantizing all the dimensions of a vector at once, we first split the vectors into subvectors of equal length, and then quantize each sub-vectors using a shared codebook. The main advantage of split quantization when applied to the VQ-VAE architecture is that one may vary the degree of spatial pooling without changing the total number of quantized vectors per representation. In practice, to maintain the number of quantized vectors while increasing spatial pooling, we simply split along the channel dimension. We observed that the global representations’ perplexity, which indicates the level of utilization of the codebook, substantially increases when split quantization is used compared to standard quantization (Fig. 1c). As shown in Supp. Fig. 1, split quantization is performed along the channel dimension by splitting each channel-wise vector into nine parts, and quantizing each of the resulting ‘sub-vectors’ against the same codebook. Split quantization is only needed for the global representation.

### Global and local representations

The dimensions of the global and local representations are 4 × 4 × 576 and 25 × 25 × 64 voxel, respectively. These two representations are quantized with two separate codebooks consisting of 2048 64-dimensional features (or codes).

### Identification pretext task

The part of our model that is tasked with identifying a held-back protein is implemented as a 2-layer perceptron built by alternatively stacking fully connected layers with 1000 hidden units and non-linear ReLU layers. The output of the classifier is a one-hot encoded vector for which each coordinate corresponds to one of the 1,311 proteins. We use categorical cross entropy as classification loss during training.

### Computational efficiency

Due to the large size of our image data (1,100,253 cropped images of dimensions 100 × 100 × 2 pixel) we recognized the need to make our architecture more efficient and thus allow for more design iterations. We opted to implement the encoder using principles from the *EfficientNet* architecture to increase computational efficiency without loosing learning capacity^51^. Specifically, we split the model of *EfficientNetB0* into two parts to make the two encoders in our model (Supp.File. 5). While we did not notice a loss of performance for the encoder, *EfficientNet* did not perform as well for decoding. Therefore, we opted to keep a standard architecture based on a stack of residual blocks for the decoder^52^

### Training protocol

The whole dataset (1,100,253 cropped images) was split into 8:1:1 into training, validation and testing data, respectively. All results shown in the figures are from testing data. We used the Adam optimizer with the initial learning rate of 0.0004. The learning rate was multiplied by 0.1 every time the validation loss did not improve for 4 epochs, and the training was terminated when the validation loss did not improve for more than 12 consecutive epochs. Images were augmented by random rotation and flipping in the training phase.

### Dimensionality reduction and clustering

Dimensionality reduction is performed using Uniform Manifold Approximation and Projection (UMAP)^53^ algorithm. We used the reference open-source python package *umap-learn* (version 0.5.0) with default values for all parameters (i.e. the Euclidean distance metric, 15 nearest neighbors, and a minimal distance of 0.1). We used AlignedUMAP for the clustering performance evaluation to facilitate the comparison of the different projections derived from all seven variants of our model (Supp. Fig. 4 and 5) or three variants of the previously described Cell inpainting model^11^ (Supp. Figs. 6 and 7). Hierarchical biclustering was performed using *seaborn* (version 0.11.1) with its default settings.

### Ground truth labels in UMAP representation

We used two sets of ground truth labels to evaluate the performance of *cytoself* at two different cellular scales, a manually curated list of proteins with exclusive organelle-level localization patterns (Supp. File 3) and 38 protein complexes collected from the CORUM database ^39^ (Supp. File 1). The 38 protein complexes were collected based on the following conditions: i) all subunits are present in the OpenCell data, ii) no overlapping subunit across the complexes, iii) each protein complex consists of more than 1 distinct subunit.

### Clustering score

To calculate a clustering score, we assume a collection of *n* points (vectors) in ℝ^*m*^: *S* = {*x*_*i*_ *∈* ℝ^*m*^ |0 ≤ *i* ≤ *n}*, and that we have a (ground truth) assignment of each point *x*_*i*_ to a class *C*_*j*_, and these classes form a partition of *S*:

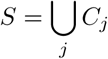

Ideally, the vectors *x*_*i*_ are such that all points in a class are tightly grouped together, and that the centroids of each class are as far apart from each other as possible. This intuition is captured in the following definition of our clustering score:

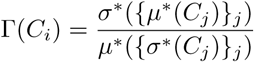

Where {.}_*k*_ denotes the set of values obtained by evaluating the expression for each value of parameter *k*, and where *µ*^∗^ and *σ*^∗^ stand for the robust mean (median) and robust standard deviation (computed using medians). Variance statistics were obtained by training the model variant 5 times followed by computing the UMAP 10 times per trained model.

### Feature spectrum

Supp. Fig. 2a illustrates the workflow for constructing the feature spectra. Specifically, we first obtain the indices of quantized vectors in the latent representation for each image crop, and then calculate the histogram of indices in all images of each protein. As a result, we obtain a matrix of histograms in which rows correspond to protein identification (ID) and columns to the feature indices (Supp. Fig. 2b). At this point, the order of the columns (that is, the feature indices) is arbitrary. Yet, different features might be highly correlated and thus either related or even redundant (depending on how “saturated” the codebook is). To meaningfully order the feature indices, we compute the Pearson correlation coefficient between the feature index “profiles” (the columns of the matrix) for each pair of feature indices to obtain a 2048 × 2048 pairwise correlation matrix (see Supp. Fig. 2c). Next we perform hierarchical biclustering in which the feature indices with the most similar profiles are iteratively merged^54^. The result is that features that have similar profiles are grouped together (Supp. Fig. 2d). This ordering yields a more meaningful and interpretable view of the whole spectrum of feature indices. We identified a number of clusters from the top levels of the feature hierarchy and manually segment them into 11 major feature clusters (ordered *i* through *xi*). Finally, for a given protein, we can produce a interpretable feature spectrum by ordering the horizontal axis of the quantized vectors histogram in the same way.

### Training cell inpainting model on OpenCell data

The cell inpainting model was constructed using the code provided by its original authors (https://github.com/alexxijielu/paired_cell_inpainting). The whole dataset was split into training, validation and testing sets (8:1:1). All results shown in the figures are computed on the basis of the test set. We used the Adam optimizer with the initial learning rate of 0.0004. The learning rate was multiplied by 0.1 every time the validation loss did not improve for 4 epochs, and the training was terminated when the validation loss did not improve for more than 12 consecutive epochs. The features to generate UMAP were extracted from layers denoted as “conv3 1”, “conv4 1” and “conv5 1” by the authors.

### Applying *cytoself* on Allen Institute dataset

Image data from the Allen Institute was downloaded from https://www.allencell.org/data-downloading.html#DownloadImageData. Patches were made following the same procedure as OpenCell dataset including max-intensity projection and downsampling to match their pixel resolutions. Nuclear center was determined using the included nuclear label in the Allen Institute dataset. We randomly selected 80 patches per protein and used these for analysis.

### Feature extraction with CellProfiler

CellProfiler 4.2.1 was used to extract features from nuclear images (without distance transform) and fluorescence protein images. In the case of *cytoself* we compute all features compatible to the data including texture features up to scale 15, for a total of 1397 features that required 2 days of computation. Only features that do not require object detection are used, including granularity, texture and the correlations between the two channels. Each feature was standardized by subtracting its mean followed by dividing by its standard deviation before a downstream analysis.

### Evaluation of the relationship between proximity in feature space and protein complex membership

The Pearson’s correlations between any two proteins in the intersection of OpenCell and CORUM database are computed with their feature spectra as the proximity metrics in the feature space. For each protein, find the ‘nearest protein’ with which it has the highest correlation, and increment the number if the correlation is higher than a given threshold, and if both of them share at least one complex in the CORUM database. To take into account the strength of correlation, we vary the minimal correlation threshold thus obtaining the curve shown in Supp. Fig. 15b.

### Statistical analysis

All box plots were generated using *matplotlib* (version 3.4.2). Each box indicates the extent from the first to the third quartile of the data, with a line representing the median. The whiskers indicates 1.5 times the inter-quartile range. *scipy* (version 1.8.0) was used to compute P values and Pearson’s correlations.

### Software and hardware

All deep learning architectures were implemented in TensorFlow 1.15^55^ on Python 3.7. Training was performed on NVIDIA V100-32GB GPUs.

## Supporting information

Supplementary files (paper v3)

## Acknowledgements

We would like to thank our colleagues at the CZ Biohub: Sandra Schmid, Mirella Bucci, Ahmet Can Solak, Bing Yang, Merlin Lange, Shruthi Vijaykumar, Luke Hyman, Marco Hein, for insightful discussions, feedback and for reviewing the manuscript. Thanks to Kibeom Kim for assistance in data analysis. Thanks to our colleague Michael Wu and James Zou from Stanford University for advice. We would like to thank Ashely Lakoduk, Jordão Bragantini and Sandra Schmid for reviewing the manuscript, and Ahmet Can Solak for helping with coding. Finally, thanks to Japan Society for the Promotion of Science and its overseas research fellowships and the *Chan Zuckerberg Biohub* and its donors for funding this work.

## Code and data availability

Source code for the models used in this work is available at: https://github.com/royerlab/cytoself

## Correspondence

Correspondence and requests for materials should be addressed to Hirofumi Kobayashi, Manuel Leonetti and Loic A. Royer

## Supplementary Text

### Interpreting the features as patterns in the images

An important and very active area of research in deep learning is the visualization, interpretation, and reverse-engineering of the inner working of deep neural networks^56,57^. To better understand the relationship between our input images and the emergent features obtained by *cytoself*, we conducted an experiment in which images were passed into the autoencoder while zeroing a given feature range before decoding. By computing the difference between the reconstructed images with or without zeroing, we identify specific regions of the images that are impacted, and thus causally linked, to that feature. Three examples are illustrated in Fig. 16: (a) POLR2E, a core subunit shared between RNA polymerases I, II and III, (b) SEC22B, a vesicle-trafficking protein, and (c) RPS18, a ribosomal protein. For each protein we highlight (in red, Supp. Fig. 16a-c) regions of the images that correspond to the three strongest peaks in their respective spectra. These difference maps reveal the image patterns that are lost and hence linked to that peak. The strongest peak (leftmost) of POLR2E’s spectrum clearly corresponds to high intensity punctate structures within nucleoli, a localization recently established by Abraham *et al*.^58^, *while the two other peaks correspond to lower intensity and more diffuse patterns. In the case of SEC22B the strongest peak (leftmost) corresponds to cytoplasmic regions with high densities of vesicles. Other peaks in the spectrum of SEC22B correspond to regions with sparse punctate expression. Finally, for RPS18, the strongest peak (rightmost) corresponds to large, diffuse, and uniform cytoplasmic regions in the images, whereas the two other selected peaks correspond to brighter and more speckled regions (middle) as well as regions adjacent to the nuclear boundary (leftmost). This analysis highlights both the interpretability but also the high complexity of the encodings generated by our model*.

### Cropping based on fiducial channel centering versus content-based centering

*Since the fiducial nuclear marker is used to centralize the input images around a nucleus it, is the marker necessary? To answer this question we trained cytoself* on a dataset cropped on the basis of the image content alone (local image entropy) – forgoing the nuclear channel entirely. We compare the clustering scores obtained from this dataset with those obtained from the dataset cropped by centering nuclei and found the difference to be negligible (see Supp. Fig. 13). This result shows that the *texture* of the protein localization patterns is more important than the relative position of the fiducial marker to the protein fluorescence, or of its position within the cropped images. The main advantage of using the nuclear fiducial marker is to optimize the layout of the crops relative to the cells. Ideally we want to have one crop per cell, and one cell per crop. In contrast, random cropping without fiducial marker cannot ensure that every cell is used.

### Dataset splitting into training, validation, and test sets

The training protocol described in the Methods section introduces data-leakage between training, validation and test data at pixel level. Another approach for splitting the data would be to split crops per field of view to ensure no that each pixel occurs only in one subset (train, validation or test). In the following we show that splitting our data along field of view does not change our results. We also explain why splitting the data into train-val-test sets is not as critical for self-supervised as it is for supervised learning.

First, we revisit our motivations for splitting the data in training, test, and validation sets. In a supervised setting, splitting the data in training, test, and validation sets serves two important purposes: (i) the test set is used to make an estimate of the performance of the model after supervised training, which is likely to generalize to further unseen data if it is *in-distribution*. the validation set is used during training to adjust the learning rate as well as to ensure early stopping to avoid over-fitting which could degrade performance on the test set. These considerations (i, ii, iii) are important for supervised learning. However, in our case, all training is self-supervised, and because the auto-encoder reconstruction and protein identification pretext *tasks* are not used *after* training and the performance metrics such as losses are not important for our end purpose. For the typical use-case of generating a feature vectors from input images, we never need to infer the identity of the tagged proteins nor do we need to reconstruct these images. While we do *not* use the pretext-tasks *per se* after training, we do use the resulting trained models and the latent representations that these models produce for given input images. Instead, we evaluated these models independently using our clustering score based on manually curated localization annotations. It follows then, that with our approach we could simply use the full dataset for training, without splitting the data. However, in general, it is often advantageous for technical purposes to do a train-val-test split to measure model convergence and detect over-fitting. The only disadvantage perhaps is that we could have trained *cytoself* on all of our data. In an abundance of caution, we use the test data for all analysis, but we could also have used the training data for the reasons explained above. Notwithstanding, it is in general preferable to avoid over-fitting, even in a self-supervised setting.

To ensure that our model does not overfit, we split our dataset *per field-of-view* and retrained the *cytoself* model. As shown in Supp. Fig. 18a, the gap between training and validation loss does not increase after about 120 epochs and 5 days of training, indicating that over-fitting does not occur. Another piece of evidence that our model did not over-fit to the training data is that our *cytoself* model actually works on images from the Allen Cell Collection (see Supp. Fig. 11). One last point is to verify that indeed our results are not sensitive to the data-splitting method. First, we check whether the results of our ablation study still hold when splitting our dataset per field of view. To check this we recomputed the clustering scores. As shown in the Supp. Fig. 18b), the relative positions among these model variants stays roughly the same. Overall these results show that the different splitting scheme does not affect the relative performance between variants of our approach. Similarly, we recompute the UMAP in Fig. 2 of our manuscript and find no difference in how well clustered the data is (see Fig. 18c). Lastly, we redid the analysis on FAM241A and reach the same conclusion (see Fig. 18d). Overall, these results show that the technical choice of data splitting does not affect our results or conclusions.

## Supplementary Table

**Table 1:**
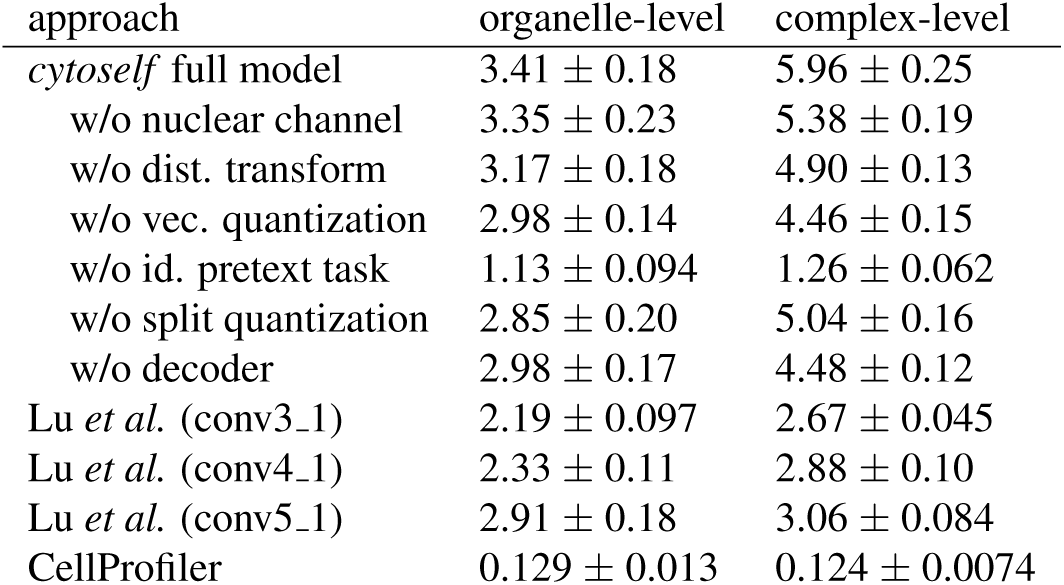
(Supplementary.) Clustering performance in our full model surpasses the previously reported cell-inpainting model^11^. We train the models 5 times, compute 10 different UMAPs, compute clustering scores using organelle-level and protein-complex-level ground truth, and then report mean and standard error of the mean (*µ* ± *sem*.). For the latent representations in the inpainting model, we examined the 3 network layers discussed in Lu *et al*. to produce image representations for UMAP. Note that our approach works with single fluorescence channel whereas the approach by Lu *et al*. needs at least two channels.

| approach | organelle-level | complex-level |
| --- | --- | --- |
| *cytoself* full model | $3.41 \pm 0.18$ | $5.96 \pm 0.25$ |
| w/o nuclear channel | $3.35 \pm 0.23$ | $5.38 \pm 0.19$ |
| w/o dist. transform | $3.17 \pm 0.18$ | $4.90 \pm 0.13$ |
| w/o vec. quantization | $2.98 \pm 0.14$ | $4.46 \pm 0.15$ |
| w/o id. pretext task | $1.13 \pm 0.094$ | $1.26 \pm 0.062$ |
| w/o split quantization | $2.85 \pm 0.20$ | $5.04 \pm 0.16$ |
| w/o decoder | $2.98 \pm 0.17$ | $4.48 \pm 0.12$ |
| Lu *et al.* (conv3_1) | $2.19 \pm 0.097$ | $2.67 \pm 0.045$ |
| Lu *et al.* (conv4_1) | $2.33 \pm 0.11$ | $2.88 \pm 0.10$ |
| Lu *et al.* (conv5_1) | $2.91 \pm 0.18$ | $3.06 \pm 0.084$ |
| CellProfiler | $0.129 \pm 0.013$ | $0.124 \pm 0.0074$ |

## Supplementary Files

1. proteins corum.csv, A list of protein subunits collected from CORUM^39^ as a ground truth to compute clustering scores. See Methods for how they were selected.

2. proteins uniloc.csv, A list of proteins that has only one localization pattern.

3. proteins uniorg.csv, A list of proteins that localizing with unique organelles.

4. proteins subunits.csv, A list of protein subunits for protein complexes mentioned in Fig. 2 and Fig. 3b.

5. model structures.zip, Detailed structure of VQ-VAE model, including **(a)** the whole model structure, **(b)** the structure of encoder1, **(c)** the structure of encoder2, **(d)** the structure of decoder1, **(e)** the structure of decoder2.

## Supplementary Figures

**Supplementary Figure 1:**
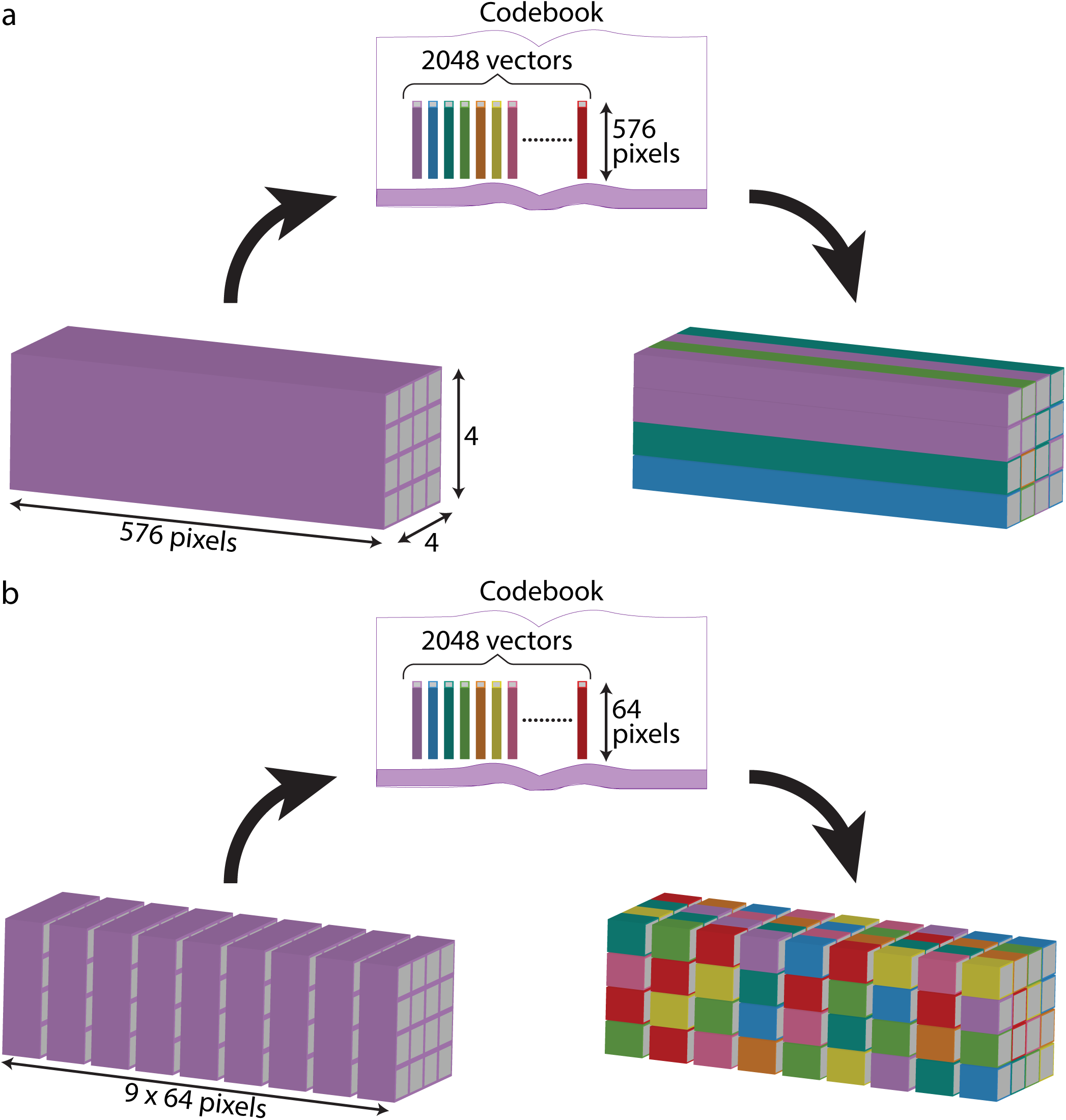
A schematic of split quantization. **(a)**, Without split quantization, there are only 4 × 4 = 16 quantized vectors in the global representation. **(b)**, With split quantization, there are 4 × 4 × 9 = 144 quantized vectors in the global representation, resulting in more opportunities for codes in the codebook to be used.

**Supplementary Figure 2:**
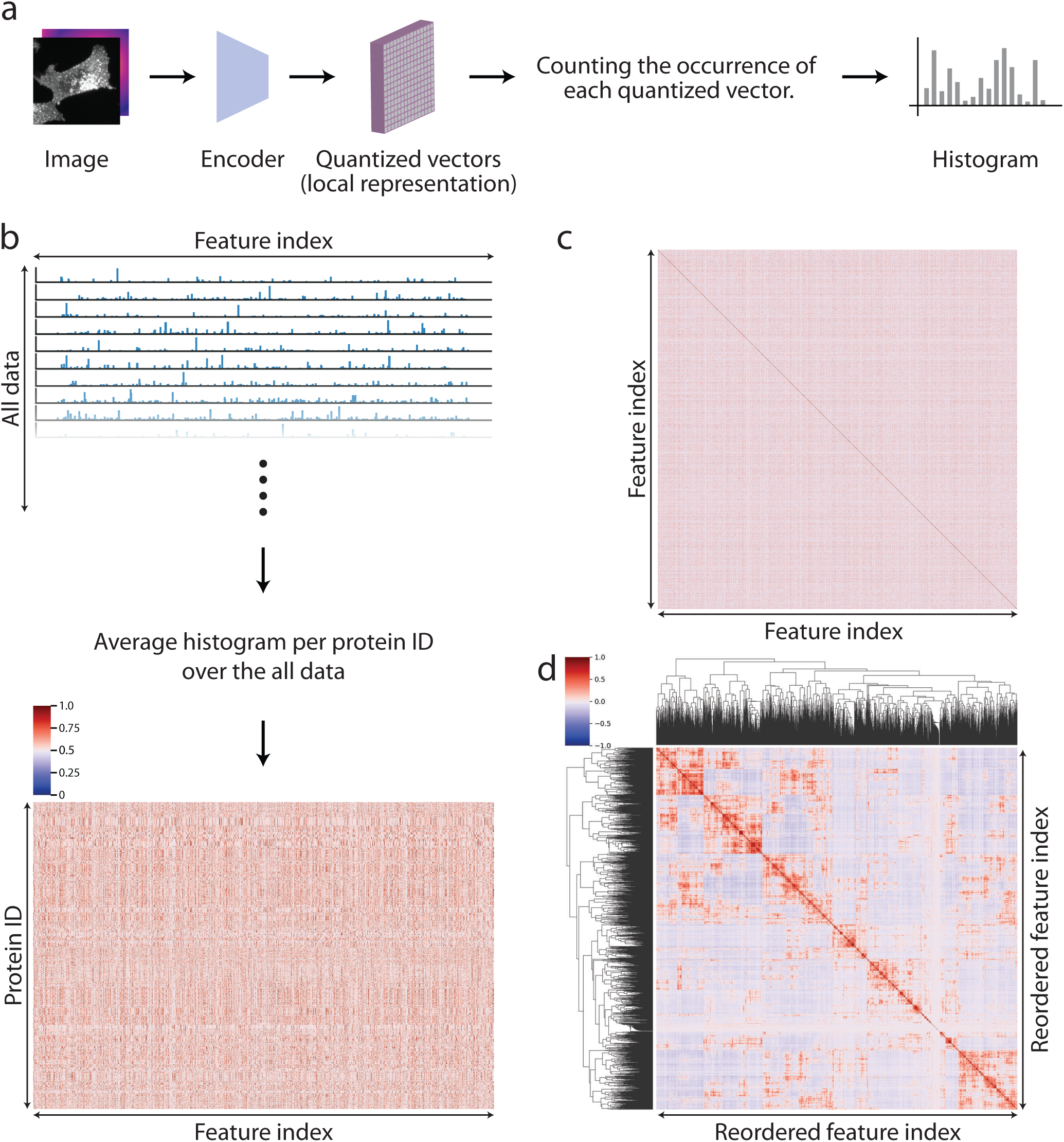
Process of constructing feature spectra. **(a)** First, the quantized vectors in the local representation were extracted and converted to a histogram by counting the occurrence of each quantized vector. **(b)** Next, taking the average of the histograms per protein ID over all the data to create a 2D histogram. **(c)** Pearson’s correlations between any two representation indices were calculated and plotted as a 2D matrix. **(d)** Finally, hierarchical clustering was performed on the correlation map so that similar features are clustered together, revealing the structure inside the local representation. The whole process corresponds to the Spectrum Conversion in Fig. 1a.

**Supplementary Figure 3:**
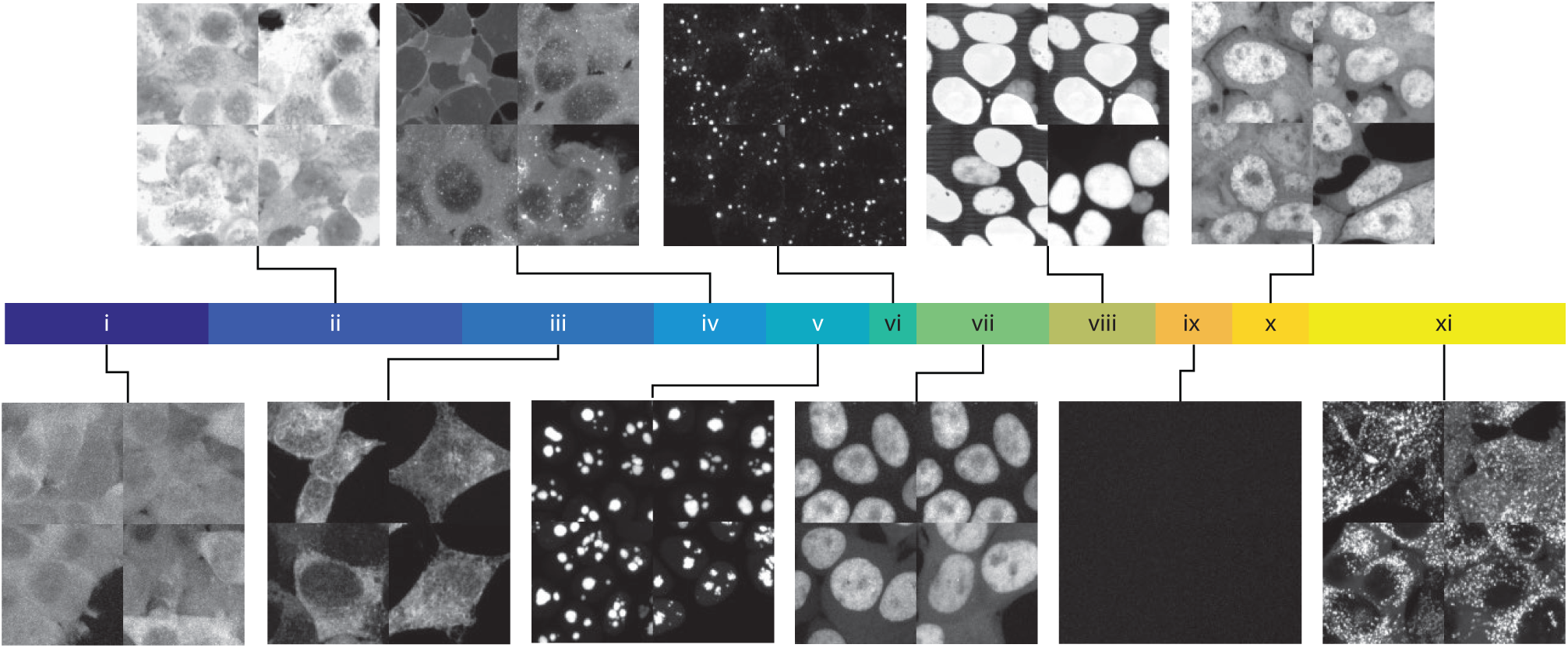
Representative images for the 11 top-level clusters. We show representative images for all 11 clusters and the corresponding localizations categories (i) cytoplasmic/membrane, (ii) cytoplasmmic/nucleoplasm, (iii) ER, (iv) membrane, (v) nucleolus, (vi) vesicles, (vii) nucleoplasm, (viii) nucleoplasm, (ix) unsuccessful image, (x) cytoplasmic/nucleoplasm, (xi) vesicles.

**Supplementary Figure 4:**
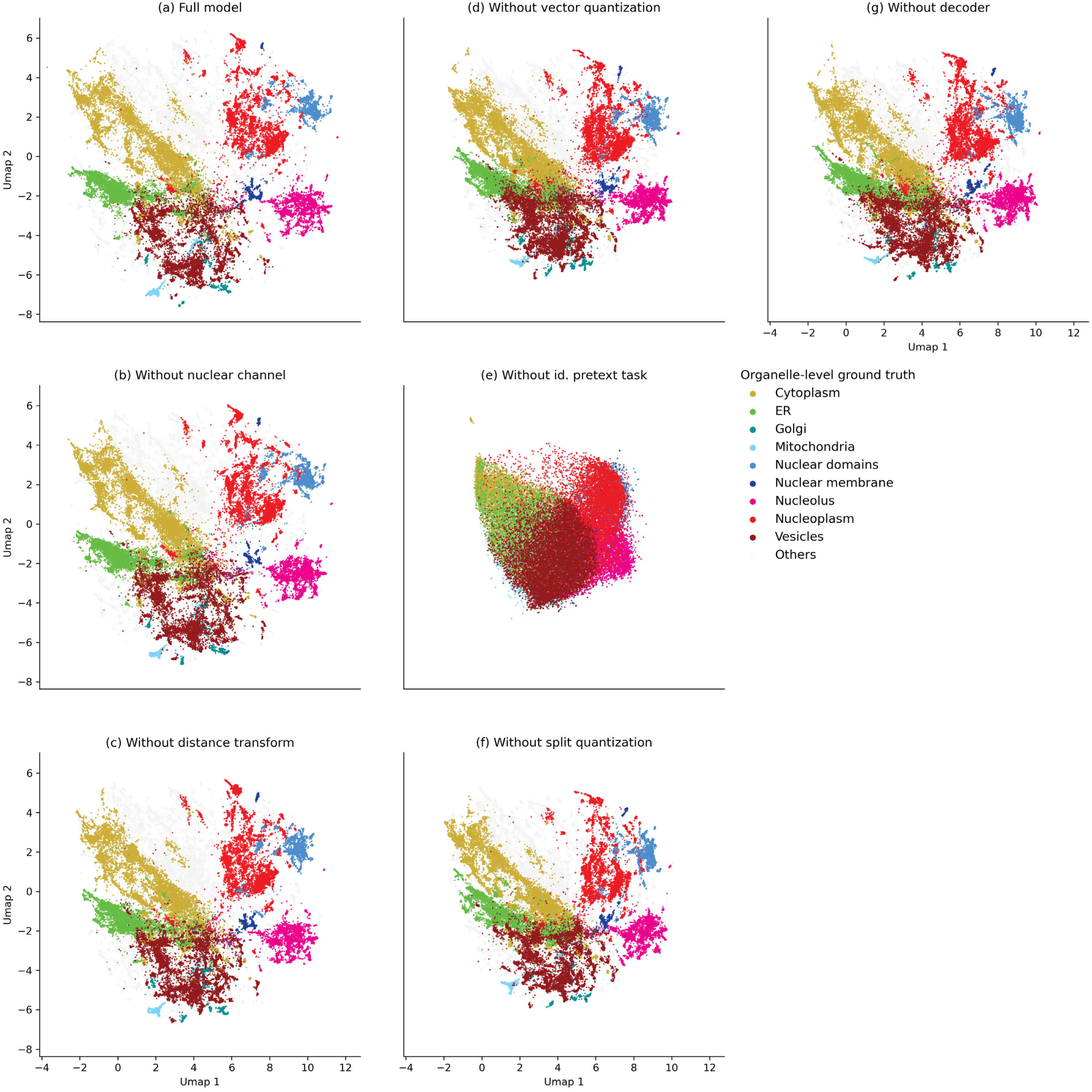
Identifying the essential components of our model with organelle-level ground truth. Protein localization UMAPs are derived after removing each components of our model separately. Aligned UMAPs are given to aid visual comparison.

**Supplementary Figure 5:**
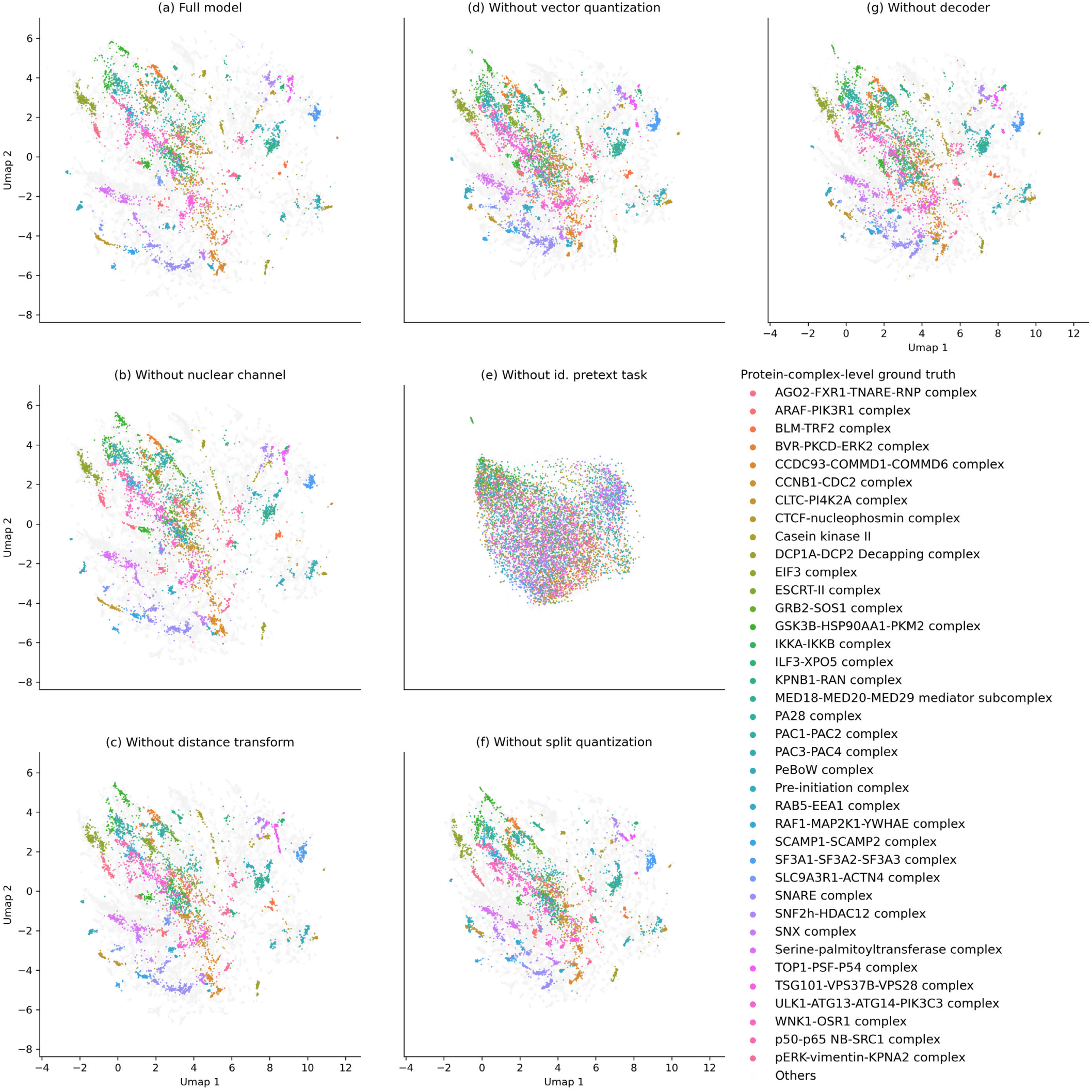
Identifying the essential components of our model with protein-complex-level ground truth. Protein localization UMAPs are derived after removing each components of our model separately. Aligned UMAPs are given to aid visual comparison.

**Supplementary Figure 6:**
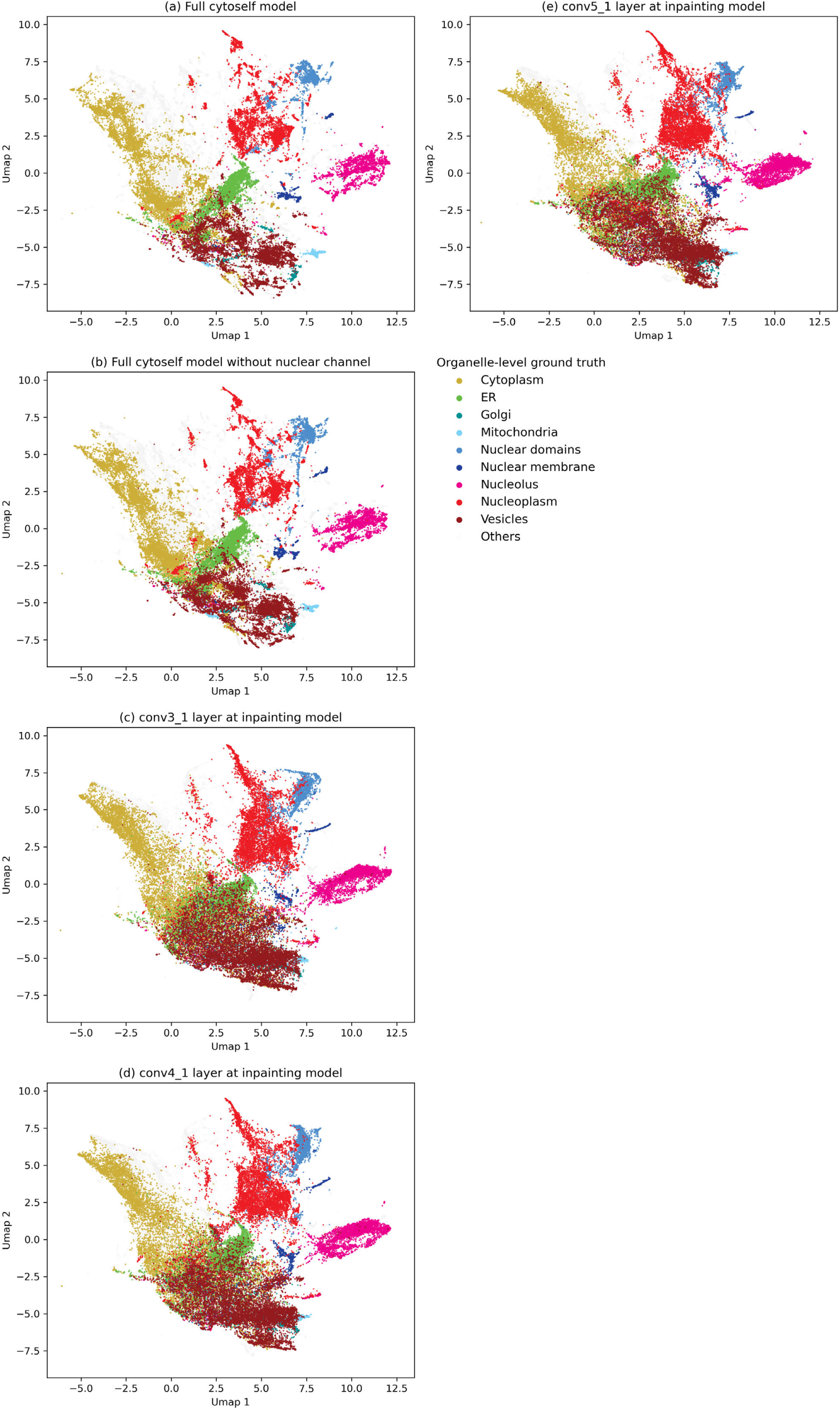
Comparing the UMAP representations between *cytoself* and cell-inpainting^11^ annotated with organelle-level ground truth. Aligned UMAPs are given to aid visual comparison.

**Supplementary Figure 7:**
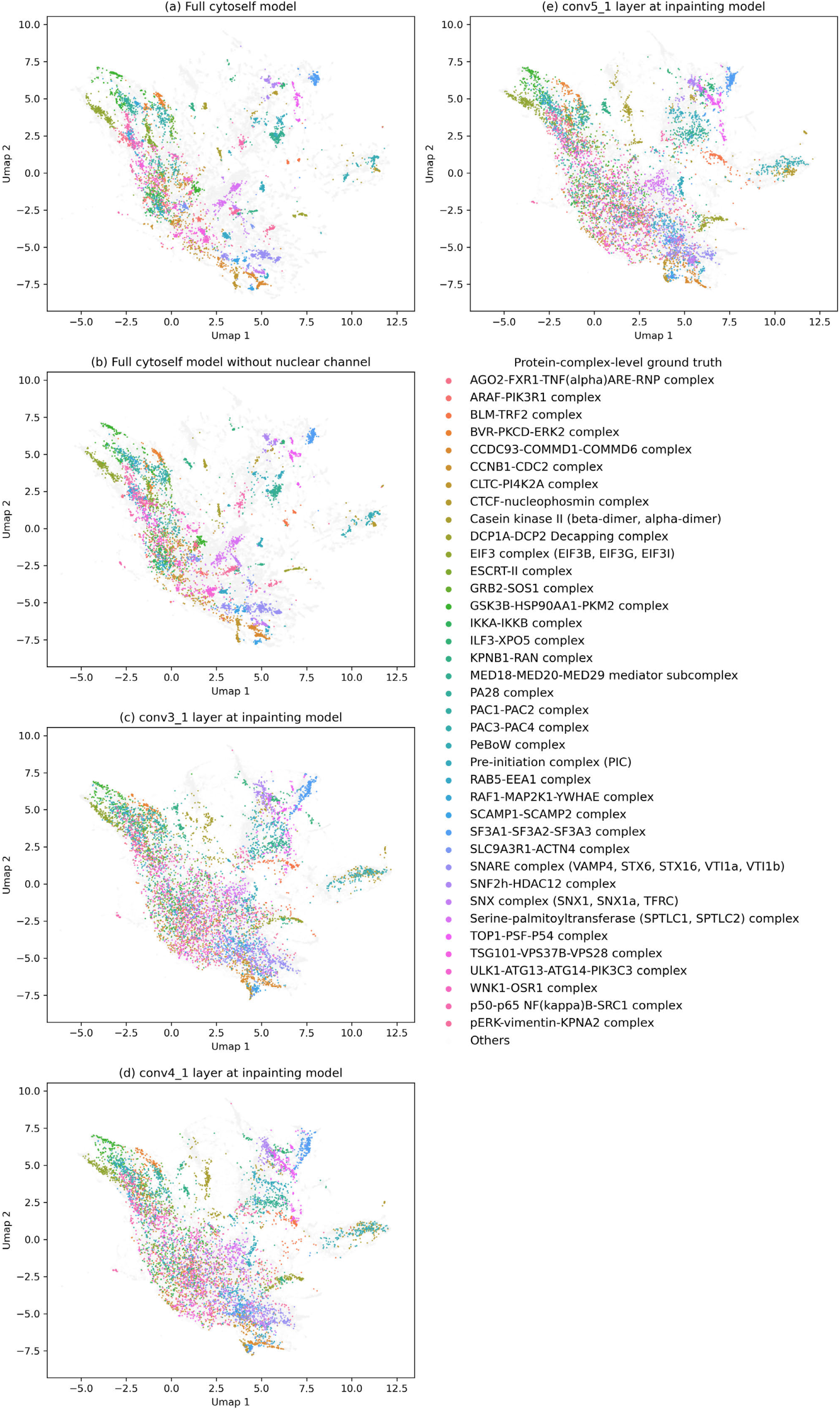
Comparing the UMAP representations between *cytoself* and cell-inpainting^11^ annotated with proteincomplex-level ground truth. Aligned UMAPs are given to aid visual comparison.

**Supplementary Figure 8:**
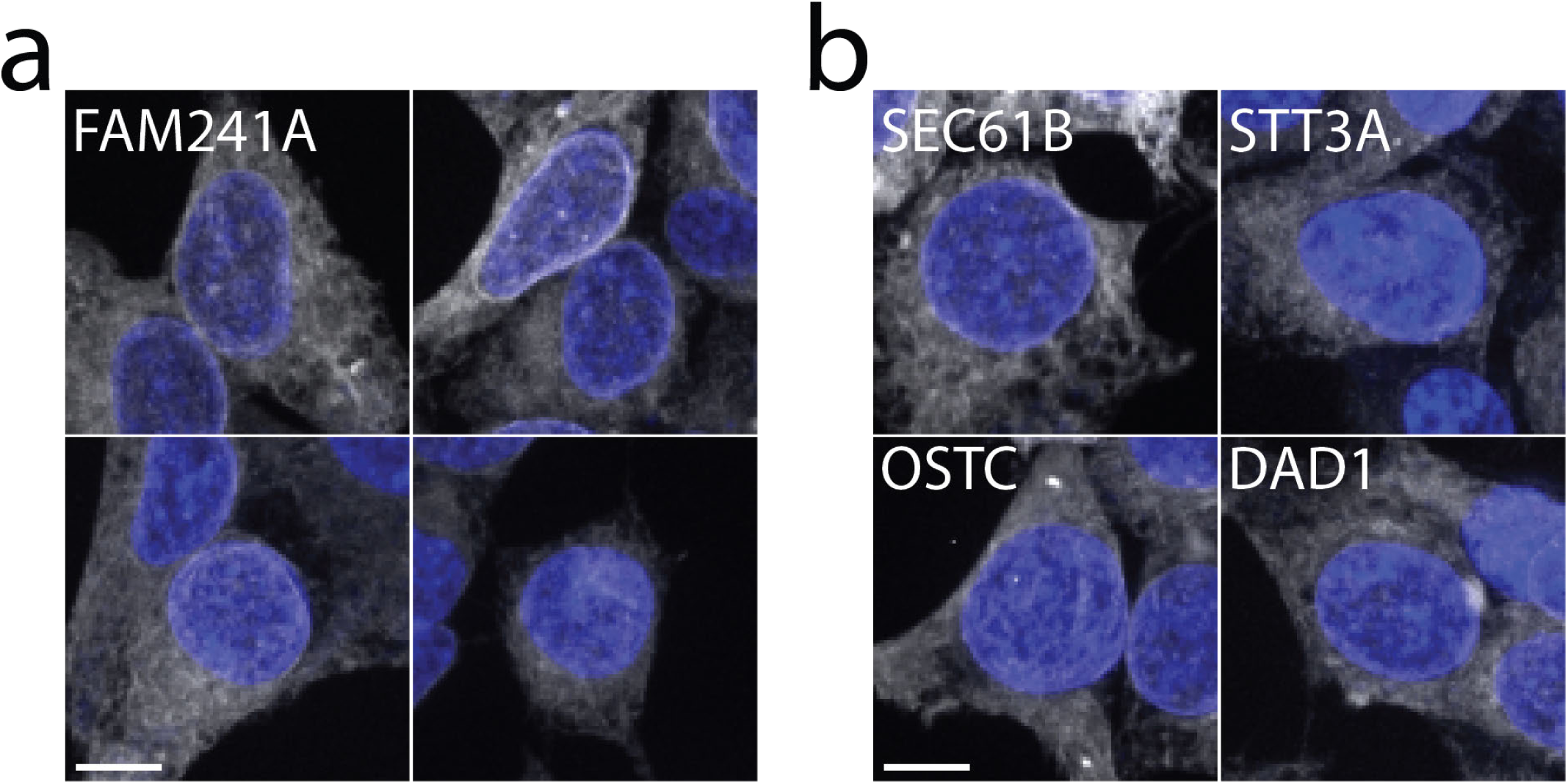
Fluorescence images for FAM241A **(a)** versus representative images of other ER localized proteins **(b)**. Protein localization and nuclei are displayed in gray and blue respectively. Scale bar: 10*µm*

**Supplementary Figure 9:**
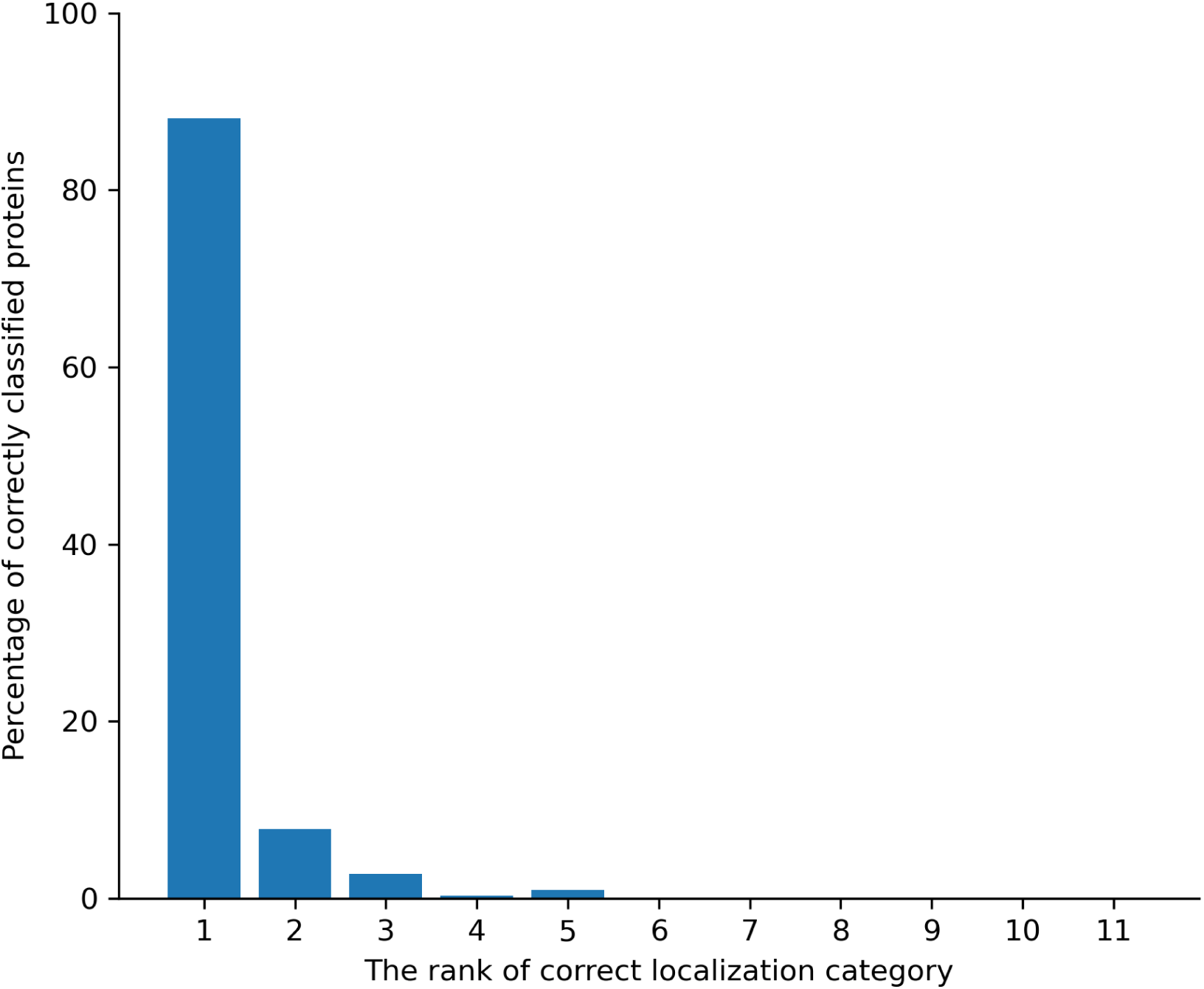
Predicting the localization category of mono-localized OpenCell proteins by correlating the *cytoself* spectra of each protein with the representative spectra of each category – in a leave-one-out fashion. Result: 88% of proteins are correctly classified. For 96% of proteins the correct annotation is within the top 2 predictions, and for 99% it is within the top 3 predictions.

**Supplementary Figure 10:**
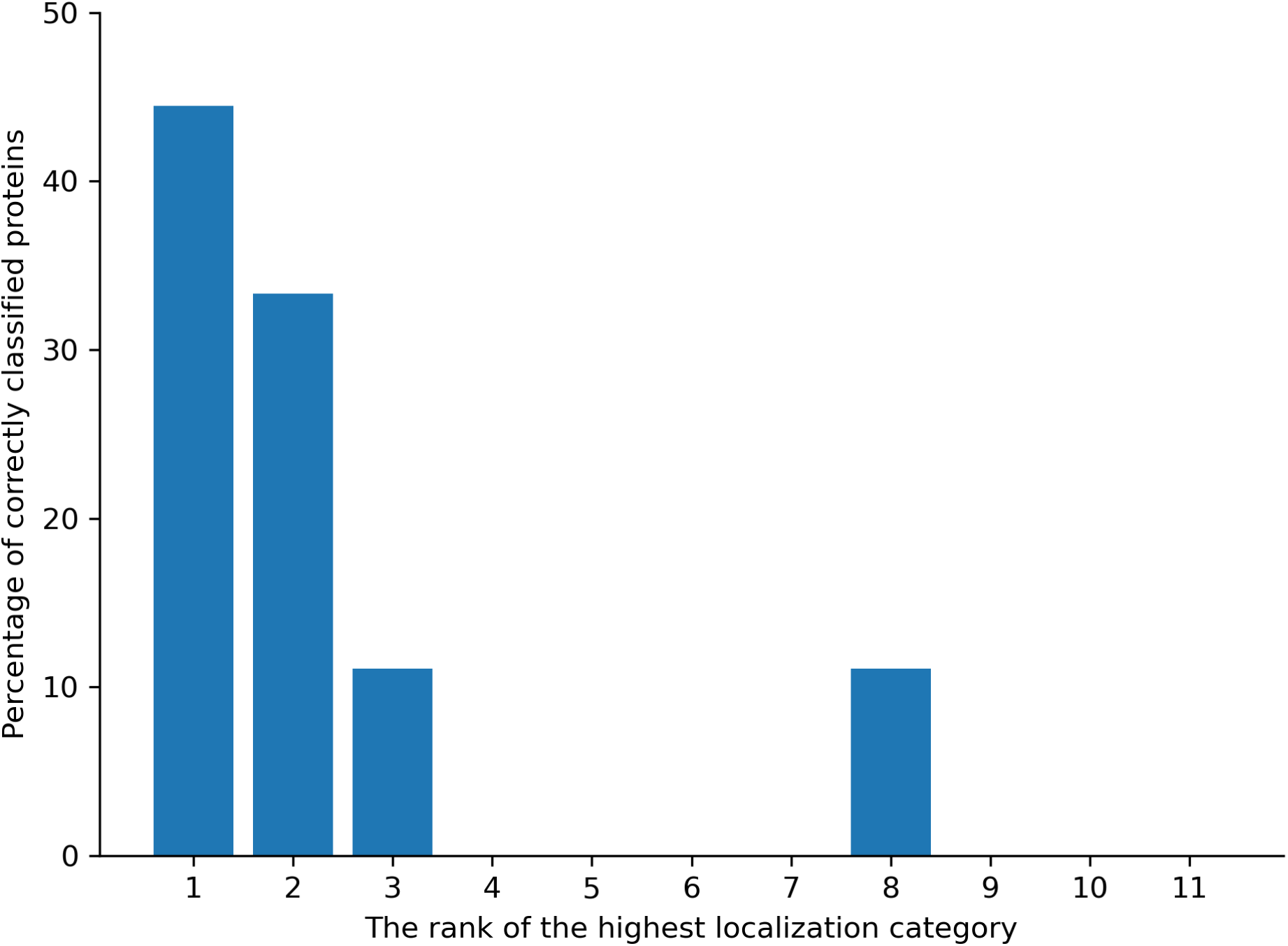
Comparing the predicted localization categories of proteins present both in OpenCell and the Allen Institute dataset. We find 9 proteins in the intersection between the two datasets (9 out of 11 in the Allen dataset). We compute the feature spectra from images from the Allen dataset, predict the corresponding localization categories, and compare these to the predictions done on the basis of the OpenCell images. Localization categories are predicted in the same way as done for FAM241A.

**Supplementary Figure 11:**
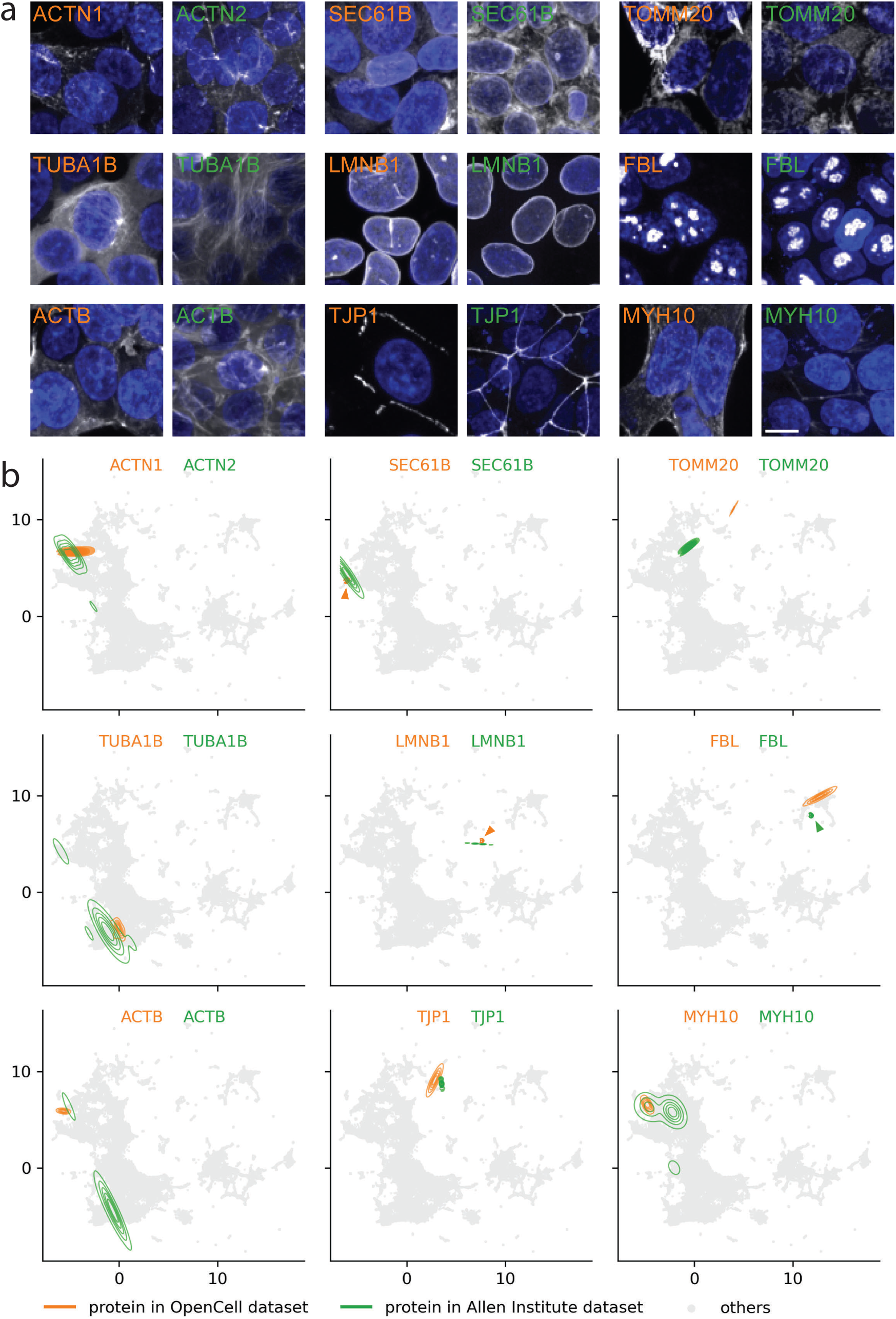
Visualizing the predicted localization categories of proteins present both in OpenCell and the Allen Institute dataset. The *cytoself* model is trained only on OpenCell data which is the same *full* model used throughout this work. Example images **(a)** and UMAPs **(b)** from OpenCell and Allen Institute datasets for the same or related proteins. The min and max intensities of each image are adjusted to ensure comparable visibility. All representative images were randomly selected. Scale bar 10 *µm*. Protein names and contour lines in orange color are from OpenCell dataset, and those in green color are from Allen Institute dataset.

**Supplementary Figure 12:**
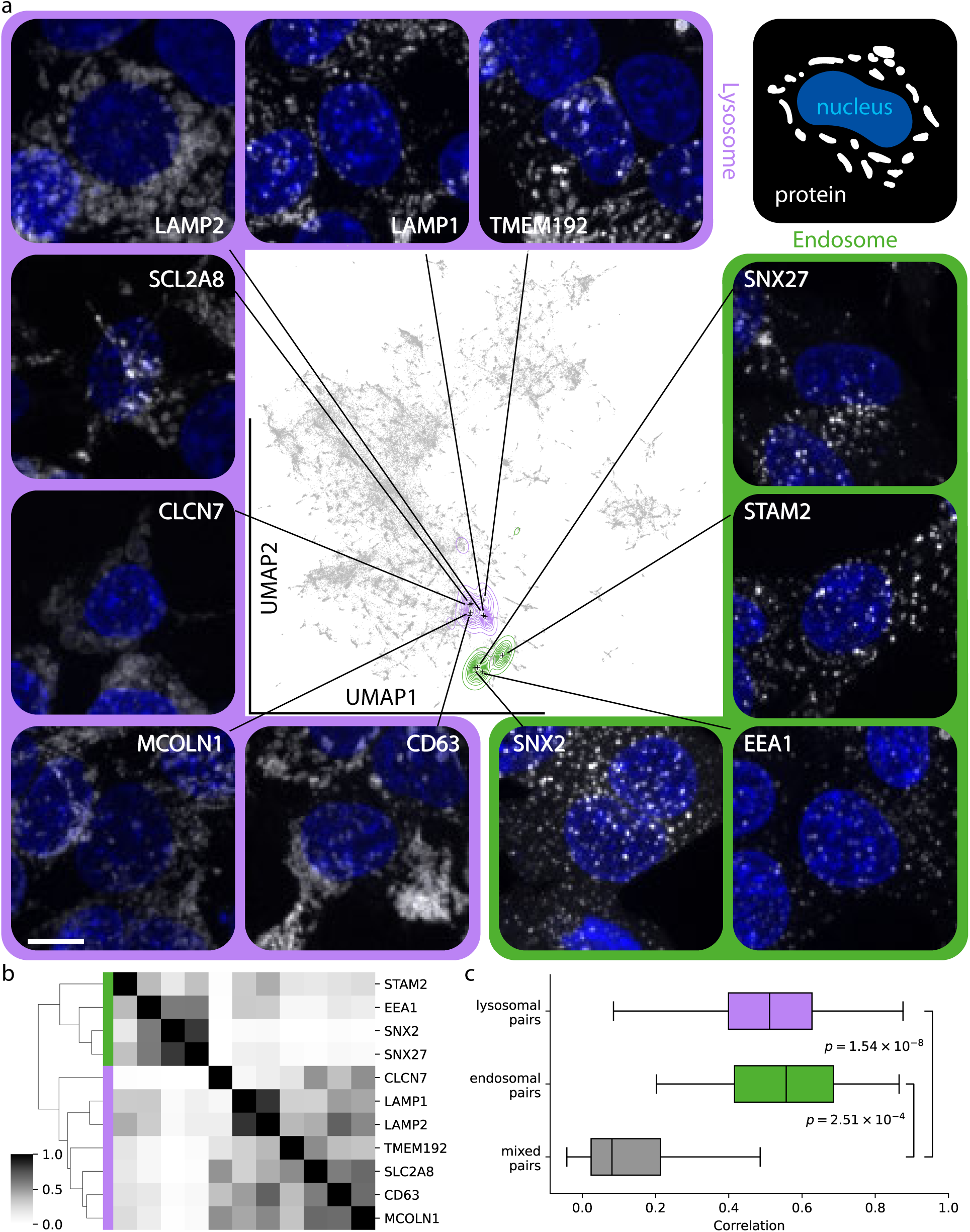
*cytoself* discriminates between lysosomal and endosomal proteins. **(a)** We selected 11 proteins in OpenCell annotated in Uniprot as lysosomal or endosomal that are independently confirmed as such by mass spectrometry^42,43^. We show that *cytoself* is able to distinguish the lysosomal from endosomal proteins solely on the basis of the fluorescence images. All of these proteins are annotated on the basis of the images as “vesicles” in both HPA and OpenCell. The min and max intensities of each image are adjusted to ensure comparable visibility. All representative images were randomly selected. Scale bar: 10 *µm*. **(b)** Clustering of these proteins on the basis of the feature spectra. **(c)** Feature spectra correlations for pairs of lysosomal and endosomal proteins, and for mixed lysosomal-endosomal pairs. The p-values are computed using the Mann-Whitney U test.

**Supplementary Figure 13:**
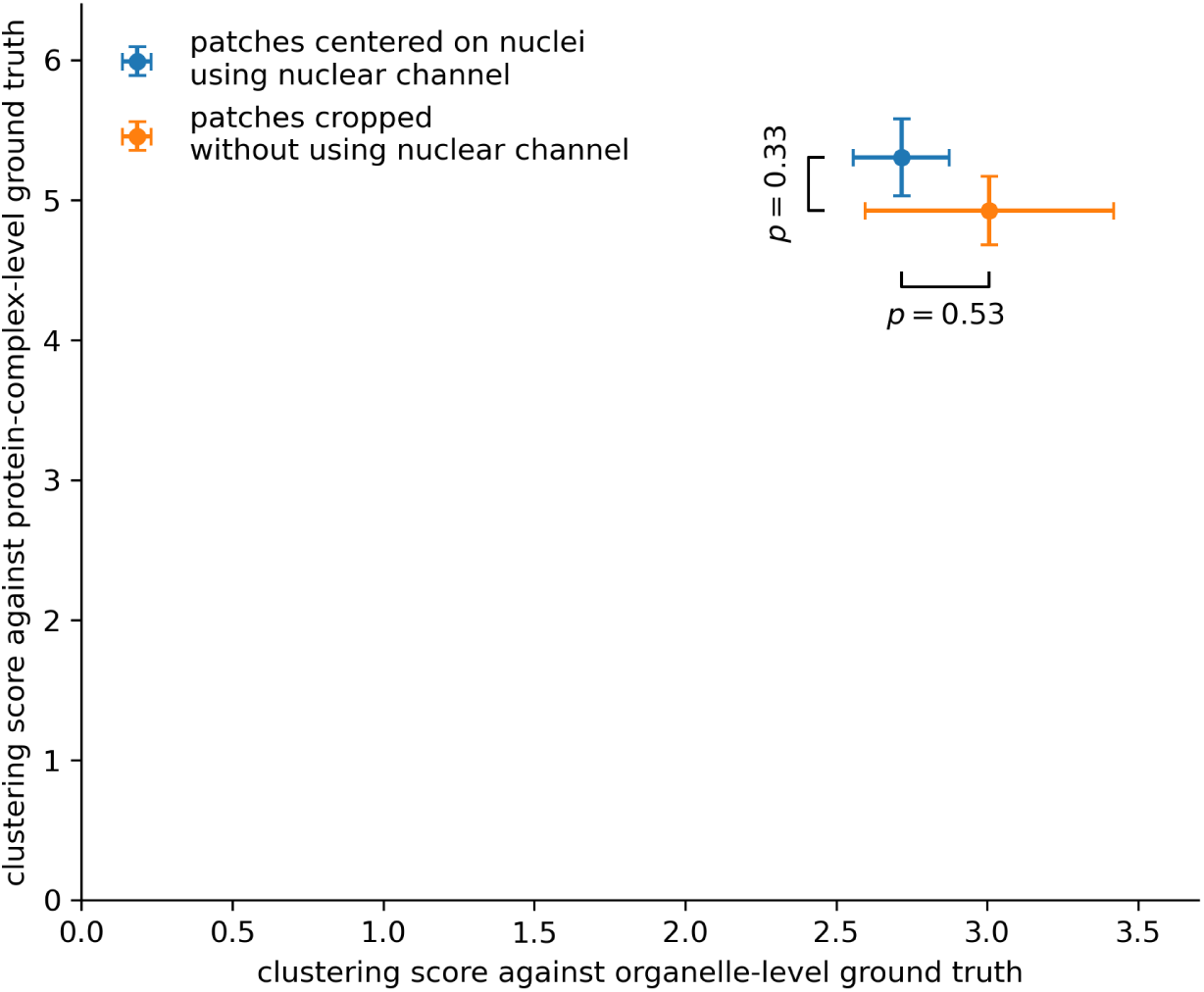
Clustering performance of *cytoself* is not significantly affected by using a training dataset cropped with or without using the nuclear fiducial channel. We avoid the use of the fiducial marker by: extracting a large number of random crops from the images, computing the histogram of each crop, computing the entropy of each of these histograms, sorting the crops by entropy, and keeping the top half of highest entropy. Variance statistics were obtained by training model variants 5 times followed by computing UMAP 10 times per trained model. The p-values are computed using Student’s t-test.

**Supplementary Figure 14:**
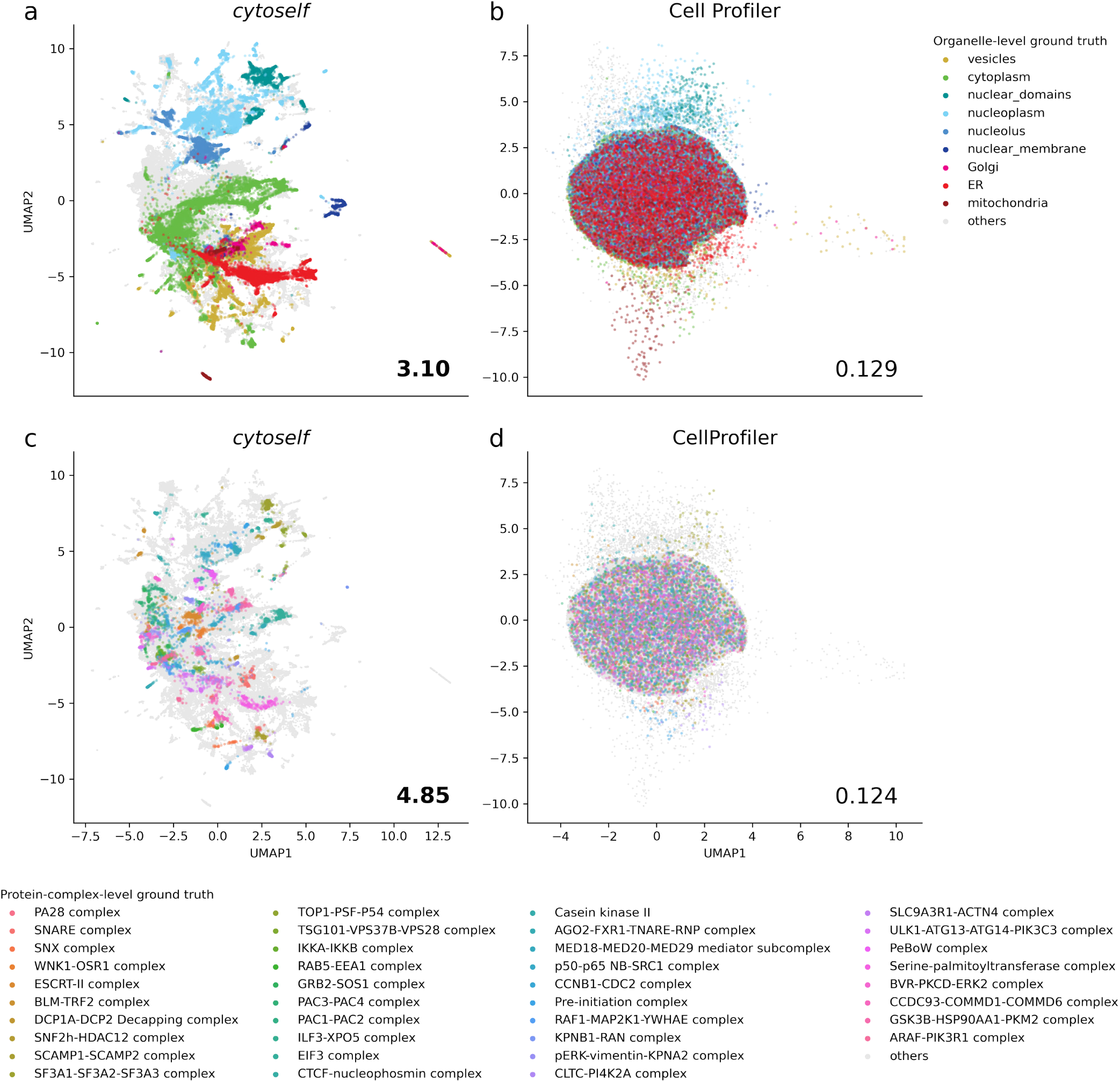
Comparing *cytoself* with CellProfiler-derived image representations. Despite our best efforts (multiple attempts with different feature normalization schemes) CellProfiler features lead to very poor clustering score (*<* 0.13) versus *cytoself* (*>* 3). **(a)** UMAP using cytoself features annotated with organelle-level ground truth. **(b)** UMAP using Cell Profiler features annotated with organelle-level ground truth. **(c)** UMAP using cytoself features annotated with protein-complex-level ground truth. **(d)** UMAP using Cell Profiler features annotated with protein-complex-level ground truth.

**Supplementary Figure 15:**
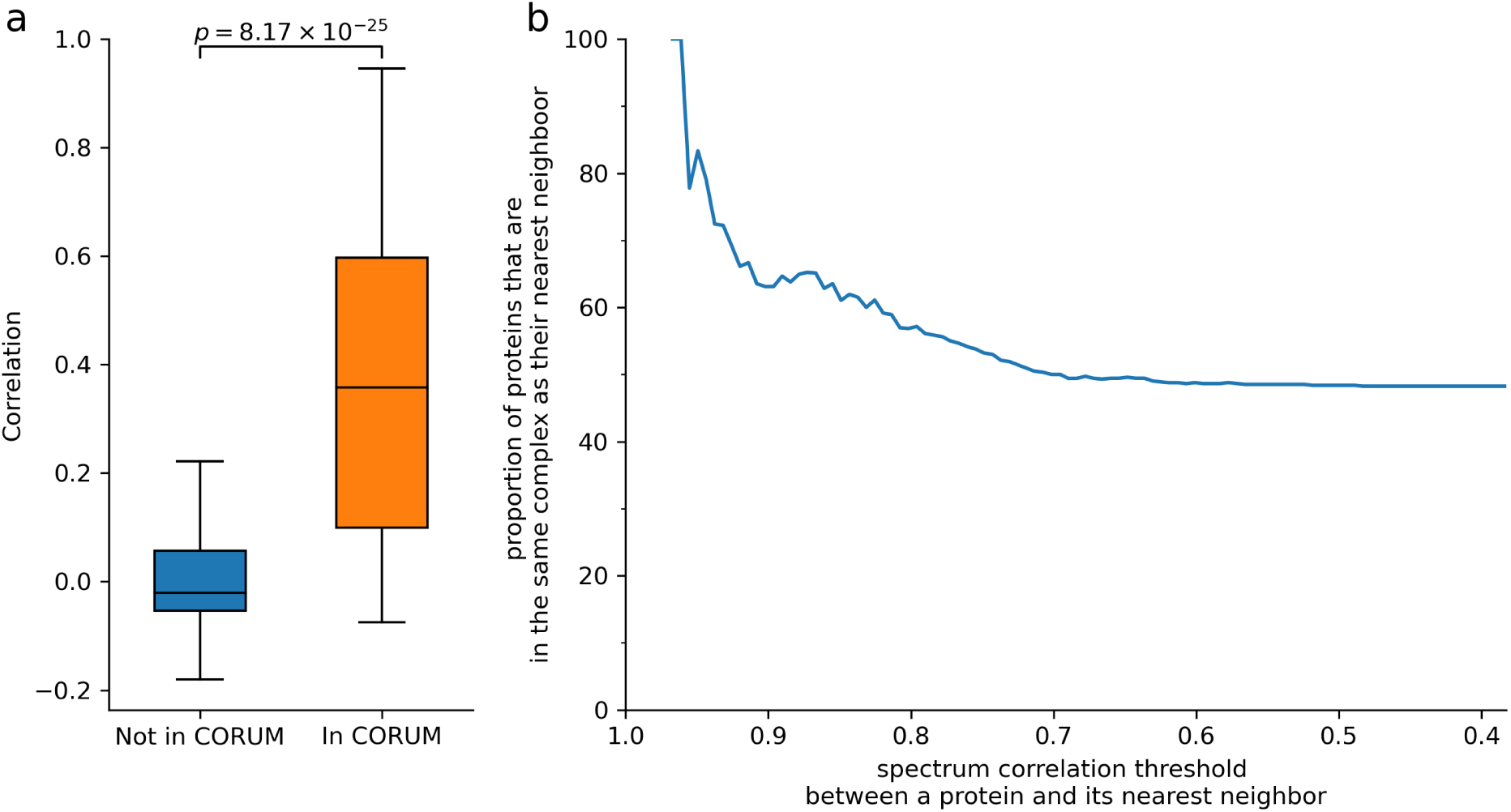
Highly correlated spectra imply shared protein complex membership. **(a)** Feature spectra of protein pairs that are in the same complex according to CORUM^39^ show significantly higher correlations than those that are not in the same complex, confirming quantitatively that the feature spectra are sensitive enough to encode complex-specific patterns. However, the spread in correlations also indicates that not all interacting proteins have strongly correlated spectra which is expected when considering that proteins can participate in different protein complexes and thus exhibit mixed localizations. In contrast, the correlation of feature spectra for protein pairs that are not in CORUM are typically close to zero with less spread, suggesting that it is rare for non-interacting proteins to have highly correlated spectra. **(b)** We plot the proportion of proteins in both OpenCell and CORUM that share protein-complex membership with their most correlated neighbor. When we consider only correlations above a threshold of 0.95 we find that in 83.3% of cases the protein with the strongest correlation is in a shared complex. For a threshold of 0.90 the value is 66.3%, and for a threshold of 0.5 the value is 47.9%. The p-value is computed using the Mann-Whitney U test.

**Supplementary Figure 16:**
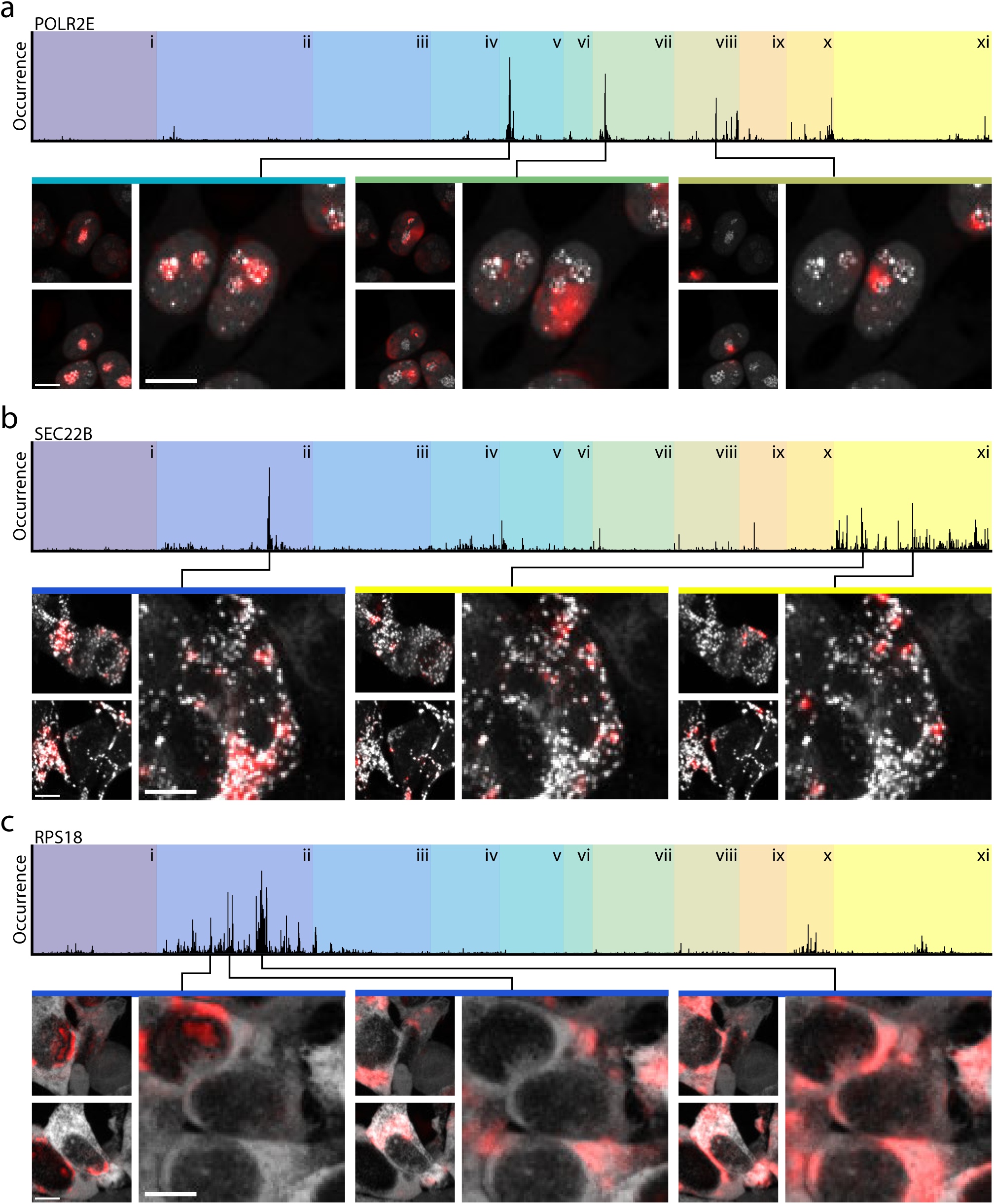
Interpreting image spectral features. Feature spectra were computed for each example proteins **(a)** POLR2E, **(b)** SEC22B, and **(c)** RPS18. Subsequently, information derived from the indicated major peaks of their feature spectra was removed by zeroing them out before passing the features again through the decoder. Highlighted in red are the differences between the resulting output images for the corresponding features and reconstructed image with full features on. The feature classes outlined in Fig. 5 are shown as background color for reference. The pixel intensities are rescaled to the minimum and maximum of each image. Scale bars: 10*µm*

**Supplementary Figure 17:**
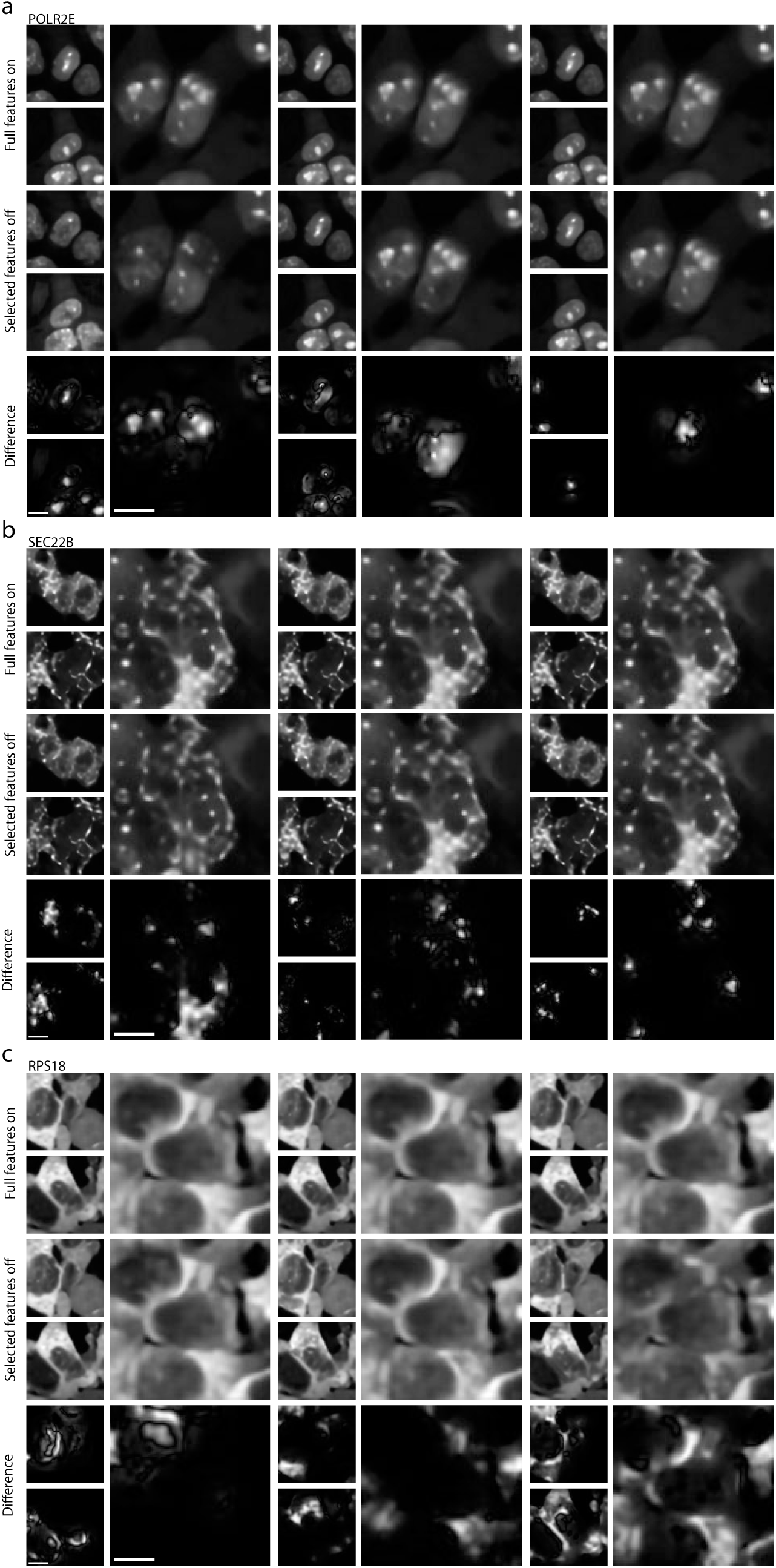
Reconstructed images with full features on, specific features off and the differences. Each panel and column correspond to those in Supp. Fig. 16. The pixel intensities are rescaled to the minimum and maximum of each image. Scale bars: 10*µm*

**Supplementary Figure 18:**
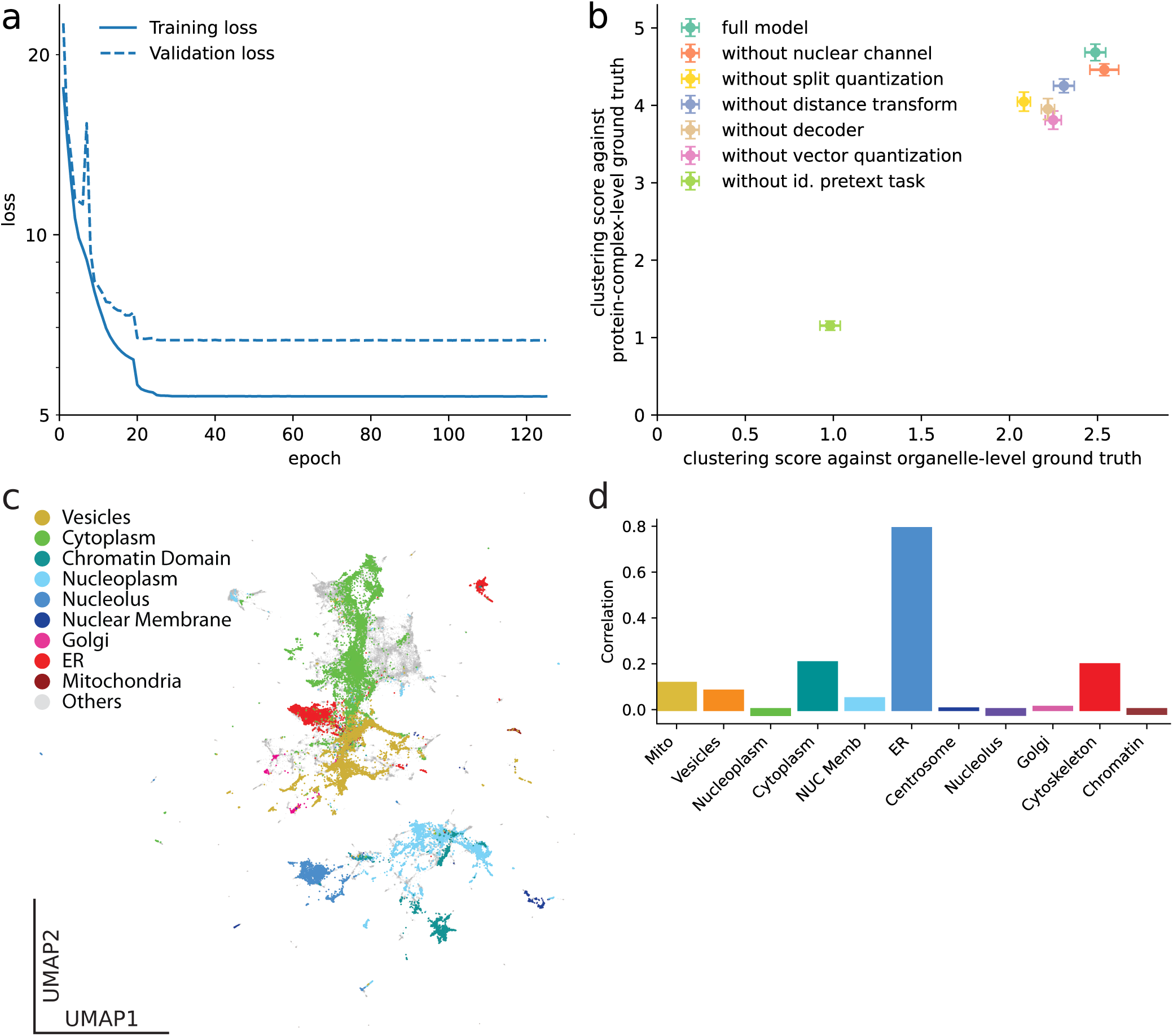
Training *cytoself* on image crops split by fields of view does not affect our results and conclusions. Splitting by fields of view ensures that each pixel occurs only once in the training data, validation data or test data, exclusively. **(a)** Training history on datasets split on fields of view showing that no over-fitting occurs even after more than 120 epochs. **(b)** Clustering scores obtained from datasets split on fields of view show that the conclusions of our ablation study are unchanged. **(c)** The appearance of the UMAP shown in Fig. 2 is not fundamentally affected by splitting on fields of view. **(d)** Splitting by fields of view does not affect our result that FAM241A is localized in the ER.

## Bibliography

1. Pepperkok, R. & Ellenberg, J. High-throughput fluorescence microscopy for systems biology. Nature reviews Molecular cell biology 7, 690–696 (2006).

2. Chandrasekaran, S. N., Ceulemans, H., Boyd, J. D. & Carpenter, A. E. Image-based profiling for drug discovery: due for a machine-learning upgrade? Nature Reviews Drug Discovery 1–15 (2020).

3. Boutros, M., Heigwer, F. & Laufer, C. Microscopy-based high-content screening. Cell 163, 1314–1325 (2015).

4. Abraham, V. C., Taylor, D. L. & Haskins, J. R. High content screening applied to large-scale cell biology. Trends in biotechnology 22, 15–22 (2004).

5. Scheeder, C., Heigwer, F. & Boutros, M. Machine learning and image-based profiling in drug discovery. Current opinion in systems biology 10, 43–52 (2018).

6. Loo, L.-H., Wu, L. F. & Altschuler, S. J. Image-based multivariate profiling of drug responses from single cells. Nature methods 4, 445–453 (2007).

7. Huh, W.-K. et al. Global analysis of protein localization in budding yeast. Nature 425, 686–691 (2003).

8. Cai, Y. et al. Experimental and computational framework for a dynamic protein atlas of human cell division. Nature 561, 411–415 (2018).

9. Thul, P. J. et al. A subcellular map of the human proteome. Science 356 (2017).

10. Cho, N. H. et al. Opencell: proteome-scale endogenous tagging enables the cartography of human cellular organization. bioRxiv (2021).

11. Lu, A. X., Kraus, O. Z., Cooper, S. & Moses, A. M. Learning unsupervised feature representations for single cell microscopy images with paired cell inpainting. PLoS computational biology 15, e1007348 (2019).

12. LeCun, Y., Bengio, Y. & Hinton, G. Deep learning. Nature 521, 436–444 (2015).

13. Perlman, Z. E. et al. Multidimensional drug profiling by automated microscopy. Science 306, 1194–1198 (2004).

14. Carpenter, A. E. et al. Cellprofiler: image analysis software for identifying and quantifying cell phenotypes. Genome biology 7, 1–11 (2006).

15. Yin, Z. et al. A screen for morphological complexity identifies regulators of switch-like transitions between discrete cell shapes. Nature cell biology 15, 860–871 (2013).

16. Bray, M.-A. et al. Cell painting, a high-content image-based assay for morphological profiling using multiplexed fluorescent dyes. Nature protocols 11, 1757 (2016).

17. Kraus, O. Z. et al. Automated analysis of high-content microscopy data with deep learning. Molecular systems biology 13, 924 (2017).

18. Eulenberg, P. et al. Reconstructing cell cycle and disease progression using deep learning. Nature Communications 8, 463 (2017).

19. Caicedo, J. C. et al. Data-analysis strategies for image-based cell profiling. Nature methods 14, 849–863 (2017).

20. Sailem, H., Bousgouni, V., Cooper, S. & Bakal, C. Cross-talk between rho and rac gtpases drives deterministic exploration of cellular shape space and morphological heterogeneity. Open biology 4, 130132 (2014).

21. Traag, V. A., Waltman, L. & Van Eck, N. J. From louvain to leiden: guaranteeing well-connected communities. Scientific reports 9, 1–12 (2019).

22. Jones, T. R. et al. Scoring diverse cellular morphologies in image-based screens with iterative feedback and machine learning. Proceedings of the National Academy of Sciences 106, 1826–1831 (2009).

23. Ouyang, W. et al. Analysis of the human protein atlas image classification competition. Nature methods 16, 1254–1261 (2019).

24. Blasi, T. et al. Label-free cell cycle analysis for highthroughput imaging flow cytometry. Nature communications 7, 1–9 (2016).

25. Pawlowski, N., Caicedo, J. C., Singh, S., Carpenter, A. E. & Storkey, A. Automating morphological profiling with generic deep convolutional networks. BioRxiv 085118 (2016).

26. Doan, M. et al. Deepometry, a framework for applying supervised and weakly supervised deep learning to imaging cytometry. Nature protocols 16, 3572–3595 (2021).

27. Goyal, P. et al. Self-supervised pretraining of visual features in the wild. arXiv preprint 2103.01988 (2021).

28. Holmberg, O. G. et al. Self-supervised retinal thickness prediction enables deep learning from unlabelled data to boost classification of diabetic retinopathy. Nature Machine Intelligence 2, 719–726 (2020).

29. Hadsell, R. et al. Learning long-range vision for autonomous off-road driving. Journal of Field Robotics 26, 120–144 (2009).

30. Batson, J. & Royer, L. Noise2self: Blind denoising by self-supervision. In International Conference on Machine Learning, 524–533 (PMLR, 2019).

31. Kobayashi, H. et al. Intelligent whole-blood imaging flow cytometry for simple, rapid, and cost-effective drugsusceptibility testing of leukemia. Lab on a Chip 19, 2688–2698 (2019).

32. Chen, T., Kornblith, S., Norouzi, M. & Hinton, G. A simple framework for contrastive learning of visual representations. In International conference on machine learning, 1597–1607 (PMLR, 2020).

33. Kolesnikov, A., Zhai, X. & Beyer, L. Revisiting selfsupervised visual representation learning. In Proceedings of the IEEE/CVF Conference on Computer Vision and Pattern Recognition, 1920–1929 (2019).

34. Deng, J. et al. Imagenet: A large-scale hierarchical image database. In 2009 IEEE conference on computer vision and pattern recognition, 248–255 (Ieee, 2009).

35. Zaritsky, A. et al. Interpretable deep learning uncovers cellular properties in label-free live cell images that are predictive of highly metastatic melanoma. Cell Systems 12, 733–747 (2021).

36. Wu, H. & Flierl, M. Vector quantization-based regularization for autoencoders. In Proceedings of the AAAI Conference on Artificial Intelligence, vol. 34, 6380–6387 (2020).

37. Van Den Oord, A., Vinyals, O. et al. Neural discrete representation learning. In Advances in Neural Information Processing Systems, 6306–6315 (2017).

38. Razavi, A., van den Oord, A. & Vinyals, O. Generating diverse high-fidelity images with vq-vae-2. In Advances in Neural Information Processing Systems, 14866–14876 (2019).

39. Giurgiu, M. et al. Corum: the comprehensive resource of mammalian protein complexes—2019. Nucleic acids research 47, D559–D563 (2019).

40. Donovan-Maiye, R. M. et al. A deep generative model of 3d single-cell organization. PLOS Computational Biology 18, e1009155 (2022).

41. Uniprot: the universal protein knowledgebase in 2021. Nucleic acids research 49, D480–D489 (2021).

42. Schröder, B. A., Wrocklage, C., Hasilik, A. & Saftig, P. The proteome of lysosomes. Proteomics 10, 4053–4076 (2010).

43. Gosney, J. A., Wilkey, D. W., Merchant, M. L. & Ceresa, B. P. Proteomics reveals novel protein associations with early endosomes in an epidermal growth factor–dependent manner. Journal of Biological Chemistry 293, 5895–5908 (2018).

44. Cheng, Y. & Church, G. M. Biclustering of expression data. In Ismb, vol. 8, 93–103 (2000).

45. Gerbin, K. A. et al. Cell states beyond transcriptomics: integrating structural organization and gene expression in hipsc-derived cardiomyocytes. Cell Systems 12, 670–687 (2021).

46. Viana, M. P. et al. Robust integrated intracellular organization of the human ips cell: where, how much, and how variable. BioRxiv 2020–12 (2021).

47. Halevy, A., Norvig, P. & Pereira, F. The unreasonable effectiveness of data. IEEE Intelligent Systems 24, 8–12 (2009).

48. Leonetti, M. D., Sekine, S., Kamiyama, D., Weissman, J. S. & Huang, B. A scalable strategy for high-throughput gfp tagging of endogenous human proteins. Proc Natl Acad Sci U S A 113, E3501–8 (2016).

49. Li, C. H. & Lee, C. Minimum cross entropy thresholding. Pattern recognition 26, 617–625 (1993).

50. Li, C. & Tam, P. K.-S. An iterative algorithm for minimum cross entropy thresholding. Pattern recognition letters 19, 771–776 (1998).

51. Tan, M. & Le, Q. Efficientnet: Rethinking model scaling for convolutional neural networks. In International Conference on Machine Learning, 6105–6114 (2019).

52. He, K., Zhang, X., Ren, S. & Sun, J. Deep residual learning for image recognition. In Proceedings of the IEEE conference on computer vision and pattern recognition, 770–778 (2016).

53. McInnes, L., Healy, J. & Melville, J. Umap: Uniform manifold approximation and projection for dimension reduction. arXiv preprint 1802.03426 (2018).

54. Rokach, L. & Maimon, O. Clustering methods. In Data mining and knowledge discovery handbook, 321–352 (Springer, 2005).

55. Abadi, M. et al. TensorFlow: Large-scale machine learning on heterogeneous systems (2015). URL https://www.tensorflow.org/. Software available from http://tensorflow.org.

56. Yosinski, J., Clune, J., Nguyen, A., Fuchs, T. & Lipson, H. Understanding neural networks through deep visualization. arXiv preprint 1506.06579 (2015).

57. Montavon, G., Samek, W. & Müller, K.-R. Methods for interpreting and understanding deep neural networks. Digital Signal Processing 73, 1–15 (2018).

58. Abraham, K. J. et al. Nucleolar rna polymerase ii drives ribosome biogenesis. Nature 585, 298–302 (2020).

