## Supplementary figures and images for "Self-Supervised Deep Learning Encodes High-Resolution Features of Protein Subcellular Localization"

### a_whole_model.png

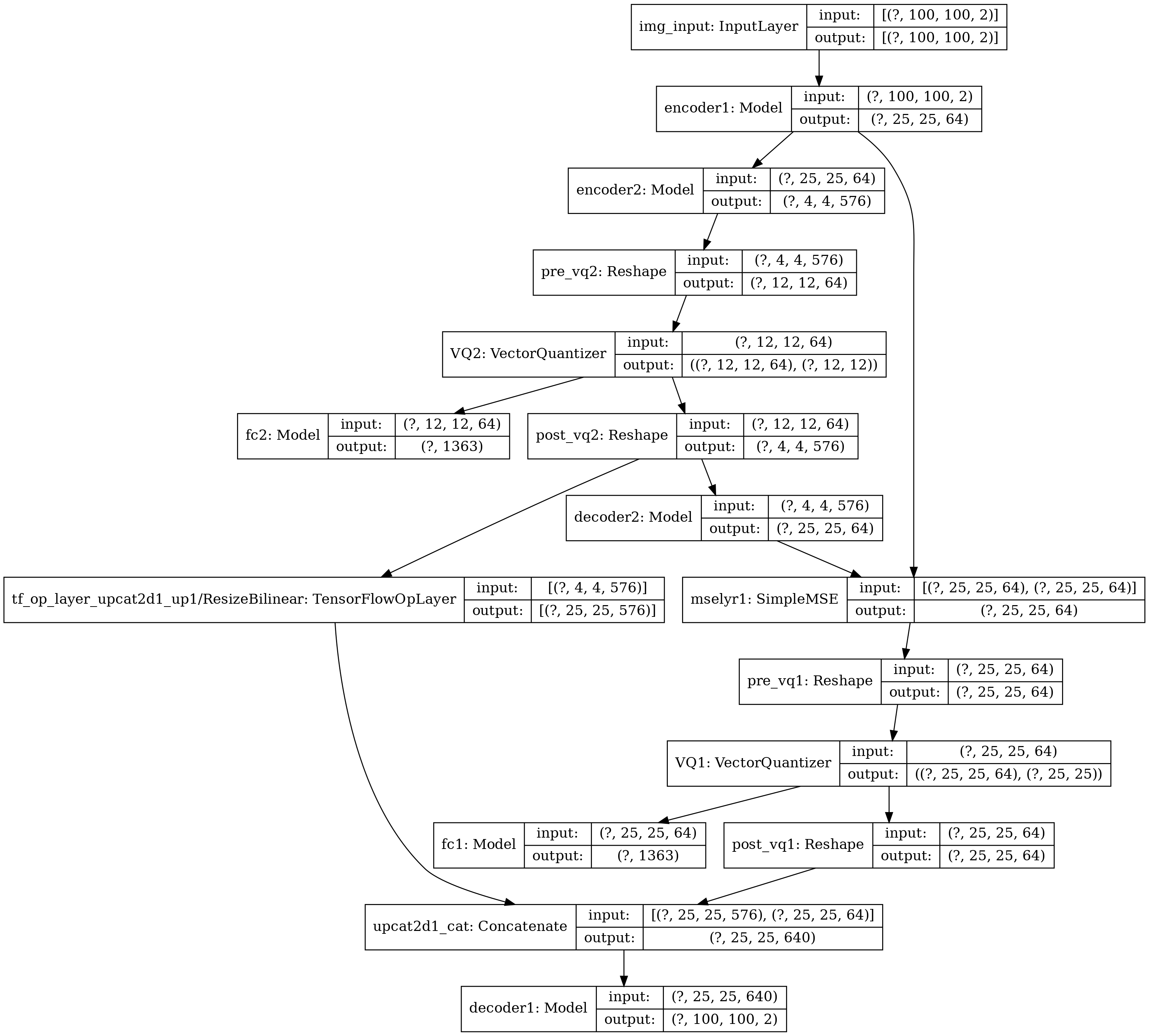

### b_encoder1.png

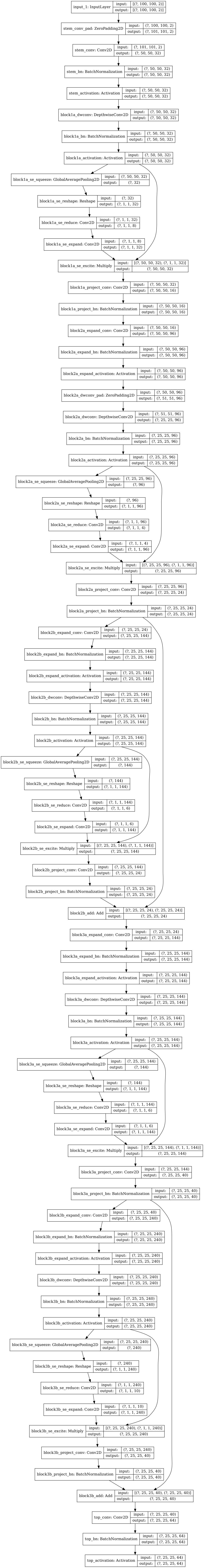

### c_encoder2.png

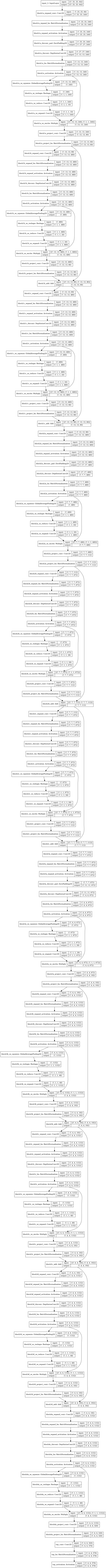

### d_decoder1.png

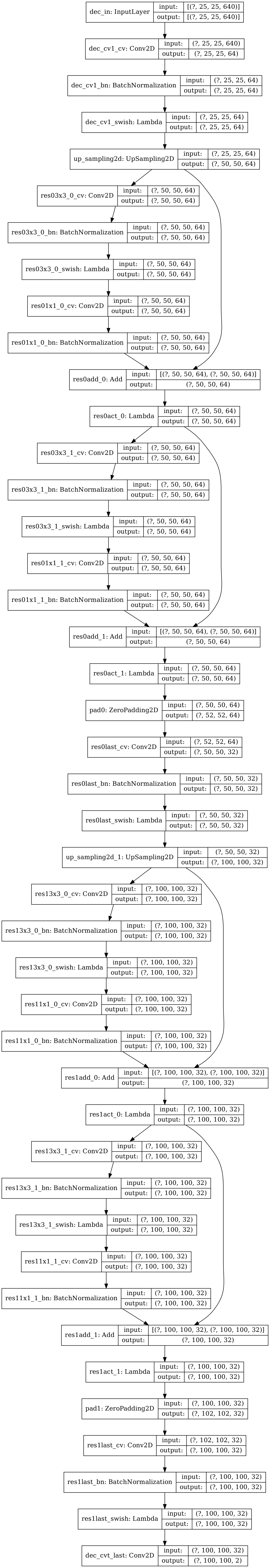

### e_decoder2.png

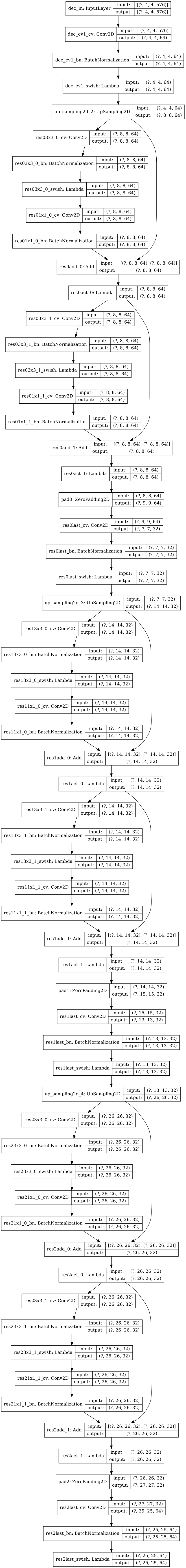
